# Cerebellum-specific deletion of the GABA_A_ receptor δ subunit alters anxiety-like, social and maternal behaviors without affecting motor performance

**DOI:** 10.1101/2019.12.27.889014

**Authors:** Stephanie Rudolph, Chong Guo, Stan Pashkovski, Tomas Osorno, Winthrop Gillis, Jeremy Krauss, Hajnalka Nyitrai, Isabella Flaquer, Mahmoud El-Rifai, Robert Sandeep Datta, Wade Regehr

## Abstract

GABA_A_ receptors containing the δGABA_A_ subunit (δGABA_A_Rs) are involved in many physiological and pathophysiological processes, such as sleep, pain, stress, anxiety-related behaviors, and postpartum depression. These extrasynaptically located, high affinity and slowly desensitizing receptors mediate tonic inhibition throughout the brain, including in granule cells (GCs) of the cerebellar input layer. However, the extent to which δGABA_A_Rs control the excitability of the cerebellar input layer and ultimately regulate behavior is unknown. We therefore deleted δGABA_A_ subunits specifically from GCs and determined the behavioral consequences in mice. Deletion reduced tonic inhibition and increased input layer excitability, but remarkably, did not affect either locomotion or motor learning. Unexpectedly, δGABA_A_ deletion heightened anxiety-like behaviors, and caused female-specific alterations in social and maternal behavior. Our findings establish that the cerebellar input layer is critical for regulating diverse behaviors that are relevant to psychiatric and neurodevelopmental disorders but were previously not associated with the cerebellum.

## Introduction

Normal function of neural circuits relies on GABAergic inhibition and its disruption can lead to disease (Ferguson and Gao, 2018; Wood et al., 2017). GABA_A_ receptor mediated inhibition occurs on different time scales and can consist of rapid (phasic) and persistent (tonic) components (Brickley et al., 1996; Farrant and Nusser, 2005; Rossi et al., 2003). Tonic inhibition influences neuronal membrane properties, such as membrane potential and resistance, and consequently controls excitability and synaptic integration (Cavelier et al., 2005; Duguid et al., 2012; Mitchell and Silver, 2003; Semyanov et al., 2004). Typically, tonic inhibition is mediated by GABA_A_ receptors containing δGABA_A_ subunits (δGABA_A_Rs) which are high affinity, slowly desensitizing, extrasynaptically located and are sensitive to ambient GABA, neurosteroids and sex hormones (Farrant and Nusser, 2005; Glykys et al., 2008; Martenson et al., 2017; Mihalek et al., 1999; Spigelman et al., 2003; Stell and Mody, 2002; Stell et al., 2003; Vicini et al., 2002; Wei et al., 2003). δGABA_A_Rs are thus poised to respond to context-dependent signals that are essential for controlling behavior. δGABA_A_Rs are involved in diverse physiological and pathophysiological processes, such as learning and memory, anxiety, stress, sleep, pain, seizures, psychiatric and neurodevelopmental disorders (Hines et al., 2012; Whissell et al., 2015), including attention deficit hyperactivity disorder (ADHD) and autism spectrum disorder (ASD) (Bridi et al., 2017; Martin et al., 2014; Zhang et al., 2017a), pregnancy and maternal behaviors (Maguire and Mody, 2008; Maguire et al., 2009), and estrous cycle-dependent fluctuations in mood and seizure susceptibility (Carver et al., 2014; Maguire et al., 2005; Smith et al., 2006). δGABA_A_Rs have therefore garnered significant interest as potential pharmacological targets for numerous disorders including postpartum depression (Melón et al., 2018; Meltzer-Brody et al., 2018), epilepsy (Petersen et al., 1983), trauma, panic and anxiety disorders (Rasmusson et al., 2017), insomnia (Orser, 2006; Thakkar et al., 2008), and some syndromic forms of ASD, such as Fragile X syndrome (FXS) (Braat and Kooy, 2015; Cogram et al., 2019; Modgil et al., 2019; Olmos-Serrano et al., 2011).

Global δGABA_A_ knock out (global δ KO) mice show diverse behavioral deficits (Maguire and Mody, 2008; Mihalek et al., 2001; Spigelman et al., 2002; Wiltgen et al., 2005), but the brain regions involved in these deficits are not known. δGABA_A_Rs are present in many brain regions, but most attention has focused on δGABA_A_s in the hippocampus, hypothalamus, amygdala and cortex, regions that have classically been associated with fear and anxiety-related behaviors, stress and cognition. In contrast, even though the cerebellum shows the highest expression levels of δGABA_A_ (Jones et al., 1997; Pirker et al., 2000; Poulter et al., 1992; Wisden et al., 1992), it has been assumed that cerebellar δGABA_A_Rs might contribute to motor learning or locomotion rather than anxiety, social behaviors or maternal behaviors (Whissell et al., 2015). However, many studies have demonstrated that the cerebellum is not limited to motor functions but controls cognitive, social and emotional processes (Buckner, 2013; Schmahmann et al., 2019; Stoodley and Schmahmann, 2018; Strick et al., 2009). In addition, animal and clinical studies found that cerebellar disruption is a common feature of psychiatric and neurodevelopmental disorders such as schizophrenia, ADHD and ASD (Becker and Stoodley, 2013; Fatemi et al., 2012; Kloth et al., 2015; Sathyanesan et al., 2019; Stoodley et al., 2017; Tsai et al., 2012; 2018). Thus, there is considerable overlap in the behavioral deficits and disorders that involve the cerebellum and δGABA_A_Rs, suggesting that cerebellar δGABA_A_Rs might contribute to diverse behaviors.

Within the cerebellar input layer, mossy fibers carry multimodal sensory information from diverse cortical, subcortical and peripheral regions and excite granule cells (GCs) (Chabrol et al., 2015; Huang et al., 2013; Ishikawa et al., 2015; Witter and De Zeeuw, 2015). Golgi cells are an important source of ambient GABA that tonically inhibits GCs by activating extrasynaptic δGABA_A_Rs that also contain the α6GABA_A_ subunit (Jechlinger et al., 1998). δGABA_A_Rs are well situated to provide an important means of regulating the excitability of the input layer. However, in previous studies, removing the α6GABA_A_ subunit eliminated tonic inhibition, but left excitability unaltered because of compensatory potassium channel expression (Brickley et al., 2001). The behavioral effects of GC hyperexcitability have therefore remained unknown. The consequences of input layer hyperexcitability are of particular interest in light of recent observations of surprisingly dense representation of GCs during certain behaviors (Badura and De Zeeuw, 2017; Cayco-Gajic and Silver, 2019; Giovannucci et al., 2017; Wagner et al., 2017), counter to classic models of the cerebellum that relied on sparse coding and a low fraction of active GCs (Marr, 1969; 1971). Thus, it is not known if elimination of δGABA_A_ subunits will alter the excitability of GCs, or be offset by compensatory mechanisms, and whether increased excitability alters behavior.

We therefore assessed whether δGABA_A_s in the cerebellum contribute to behavior. We find that cerebellar GC-specific deletion of δGABA_A_ (cb δ KO mice) decreases tonic inhibition and renders GCs hyperexcitable. Despite these profound changes to the input layer of the cerebellum, motor coordination and motor learning were unaffected in cb δ KO mice. Remarkably, cb δ KO mice displayed deficits in diverse behaviors that were previously not associated with the cerebellum, including increased anxiety-like behaviors, hyperactivity, reduced sociability in cb δ KO females, as well as abnormal maternal behavior.

## Results

Investigating the role of δGABA_A_-containing GABA_A_Rs in the cerebellum can be achieved by the selective deletion of the δGABA_A_ from cerebellar GCs. We assessed whether the Gabra6-Cre mouse line is suitable for this purpose by comparing the overlap of Cre and δGABA_A_ expression using fluorescent in situ hybridization (FISH) in whole brain sagittal sections of Gabra6-Cre x flox *tdTomato* mice (Ai14 reporter line, Figure 1A, left panels). FISH-labeled *tdTomato* RNA indicates Cre expression in the cerebellar GC layer and in the pons (Figure 1A, top), consistent with earlier descriptions of this mouse line (Fünfschilling and Reichardt, 2002). In addition, confocal images show that *tdTomato* transcripts were absent in the thalamus, dentate gyrus and the hypothalamus (Figure 1A, top). FISH indicated δGABA_A_ expression in the neocortex, striatum, dentate gyrus, thalamus, hypothalamus, and the cerebellum, but not in the pons (Figure 1A, middle), consistent with earlier reports. Thus, Cre and δGABA_A_ expression overlap selectively in cerebellar GCs (merged image, Figure 1A, bottom). We bred Gabra6-Cre mice to floxed *Gabrd* mice (Lee and Maguire, 2013) (Gabra6-Cre+, *Gabrd* flox/flox) to selectively eliminate δGABA_A_s from cerebellar GCs (cb δ KO mice).

**Figure 1:**
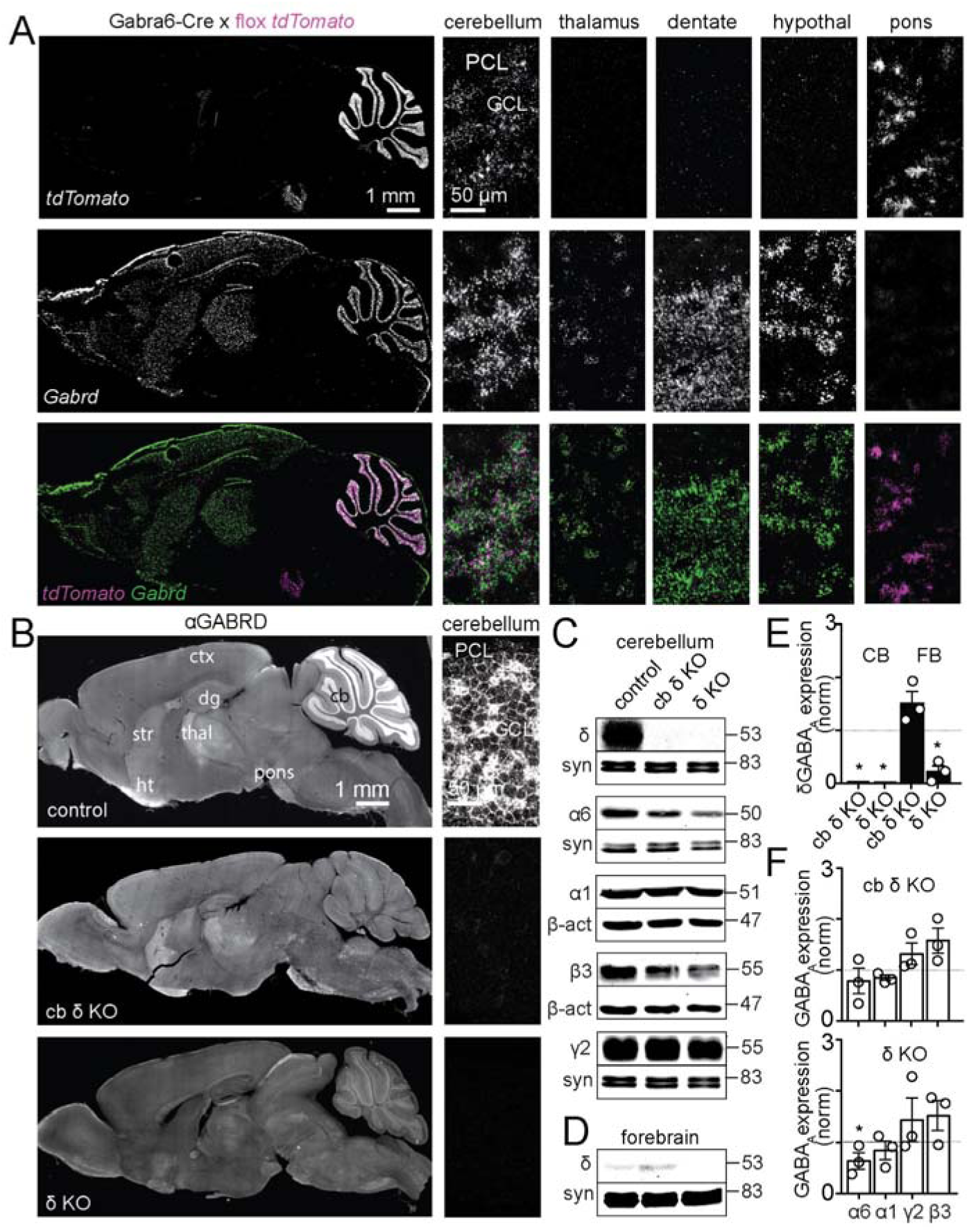
Specific deletion of the GABAA receptor δ subunit in cerebellar granule cells. A) FISH labeling of *tdTomato* transcripts in the Gabra6-cre mouse line crossed to the Ai14 reporter line (*top*), *Gabrd* transcripts *(middle*, *Gabrd*), and merged images (*bottom*) are shown. Left, sagittal section of the whole brain. Right, confocal images of the cerebellum, thalamus, dentate gyrus, hypothalamus and the pons. *tdTomato* and *Gabrd* transcripts are co-expressed exclusively in cerebellar granule cells. B) Immunostaining against the δGABA_A_ in sagittal sections of the brain (*left*) and confocal images of the cerebellum (*right*) in control (*top*), cb δ KO (*middle*), and global δ KO tissue (*bottom*). C) Western blot of the cerebellum against the δGABA_A_, α6GABA_A_, α1GABA_A_, β3GABA_A_ and γ2GABA_A_ subunits, loading controls were synapsin (syn) or β-actin (*bottom lanes*). D) Western blot of the forebrain against δGABA_A_R. E) Quantification of δGABA_A_ expression levels in the cerebellum (*white bars*) and the forebrain (*black bars*) in cb δ KO (left) and global δ KO mice. δGABA_A_ expression is abolished in the cerebellum of cb δ KO and δ KO mice and in the forebrain of δ KO mice but not in the forebrain of cb δ KO mice. (one-sample t-test of samples normalized to control, n=3 animals, p<0.05). F) Quantification of α6GABA_A_, α1GABA_A_, β3GABA_A_ and γ2GABA_A_ expression levels in the cerebellum (*white bars*) of cb δ KO mice (*middle*) and global δ KO mice (*bottom*). α6GABA_A_ expression was significantly decreased in the cerebellum of δ KO mice (p<0.05). All other receptors did not show significant changes in expression levels (one-sample t-test of samples normalized to control, n=3 animals, p>0.05)

Immunohistochemistry of parasagittal whole brain sections reveals that cerebellar GCs strongly express δGABA_A_ subunits. In control mice (*Gabrd* flox/flox), δGABA_A_ staining is the brightest in the GC layer of the cerebellum (Figure. 1 B, top left), and in confocal images of the cerebellar GC layer fluorescence is consistent with GC membranes (Figure 1 B, top right). However, in cb δ KO mice the δGABA_A_ signal was abolished almost entirely in the cerebellum (Figure 1 B, middle). An image of the background fluorescence present in global δ KO mice is shown for comparison (Figure 1 B, bottom). A comparison of the images in Figure 1 B suggests that in cb δ KO mice the δGABA_A_ subunit is eliminated from cerebellar GCs but remains intact in other brain regions. Fluorescent Western blot analysis revealed that δGABA_A_ is expressed at high levels in the cerebellum of control mice, but eliminated from cb δ KO and global δ KO animals (Figure 1 C, top panel, 1E). δGABA_A_ is expressed at relatively low levels in the forebrain of control mice, which were reduced to even lower levels in δ KO animals, while levels were slightly elevated in cb δ KO mice (Figure 1 D, top panel, 1 E). We conclude that in Gabra6-Cre x *Gabrd* flox mice, δGABA_A_ subunits are eliminated selectively from cerebellar GCs.

GABA_A_ receptor subunit deletion can affect expression of other GABA_A_ receptor subunits and, thereby, the relative contributions of phasic and tonic inhibition (Korpi et al., 2002; Nusser et al., 1999; Ogris et al., 2006; Peng et al., 2014). In cerebellar GCs tonic currents are mediated by extrasynaptic α6/δ-containing receptors, and phasic currents are mediated by α1/β3/γ2-containing subunits (Nusser et al., 1998). Besides a small reduction in α6 levels that is only significant in global δ KO mice, there were no significant differences in cerebellar α1, β3, and γ2 subunit levels in control, cb δ KO and global δ KO mice (Figure 1 C, F), suggesting that phasic currents are likely not upregulated. These findings suggest that the elimination of δGABA_A_ subunits does not have a major influence on other GABA_A_ subunits in the cerebellum.

We assessed tonic currents and the passive properties in control and cb δ KO GCs using patch clamp recordings in cerebellar brain slices (Figure 2 A-E). In control mice, a tonic current was present that increased in amplitude in the presence of 4,5,6,7-tetrahydroisoxazolo (5,4-*c*)pyridin-3(-ol) (THIP), a δGABA_A_-preferring GABA_A_ receptor agonist (Chandra et al., 2006; Meera et al., 2011; Stórustovu and Ebert, 2006), and was eliminated by blocking GABA_A_Rs with the non-selective blocker SR95531 (Figure 2 A-C, black). The initial tonic current and the increase evoked by THIP were much smaller in cb δ KO GCs (Figure 2 A-C, blue). The remaining effect of THIP is consistent with activation of non-δGABA_A_-containing receptors (Meera et al., 2011). Tonic currents were accompanied by a characteristic increase in GABA_A_R channel noise (Figure 2 A, insets, Figure S1 F). Initial tonic currents were larger in control compared to cb δ KO GCs (Figure 2 B, control, n=23; cb δ KO, n=14, p<0.0005, KS test). Tonic GABA_A_ currents in the presence of THIP were also much larger in control than in cb δ KO GCs (Figure 2 C, control, n=23; cb δ KO, n=14, p<0.0001, KS test). There was a small but significant depolarization of cb δ KO GCs (Figure 2 D, control: -61 ± 1 mV, n=28; cb δ KO: -57 ± 1 mV, n=31, p<0.05, KS test, Figure S1 B), as measured with a method that does not alter intracellular ion levels (Figure S1 A). The input resistance (R_i_) was higher in cb δ KO GCs compared to control, (Figure 2 E, control, n=18; cb δ KO, n=14, p<0.02, KS test), consistent with reduced tonic current. Additionally, in control animals, THIP decreased R_i_, while blocking GABA_A_Rs increased R_i_ (Figure S1 D), but THIP and SR95531 did not alter R_i_ in cb δ KO GCs (Figure S1 E). Thus, cb δ KO GCs show decreased tonic inhibition, resulting in increased R_i_.

**Figure 2:**
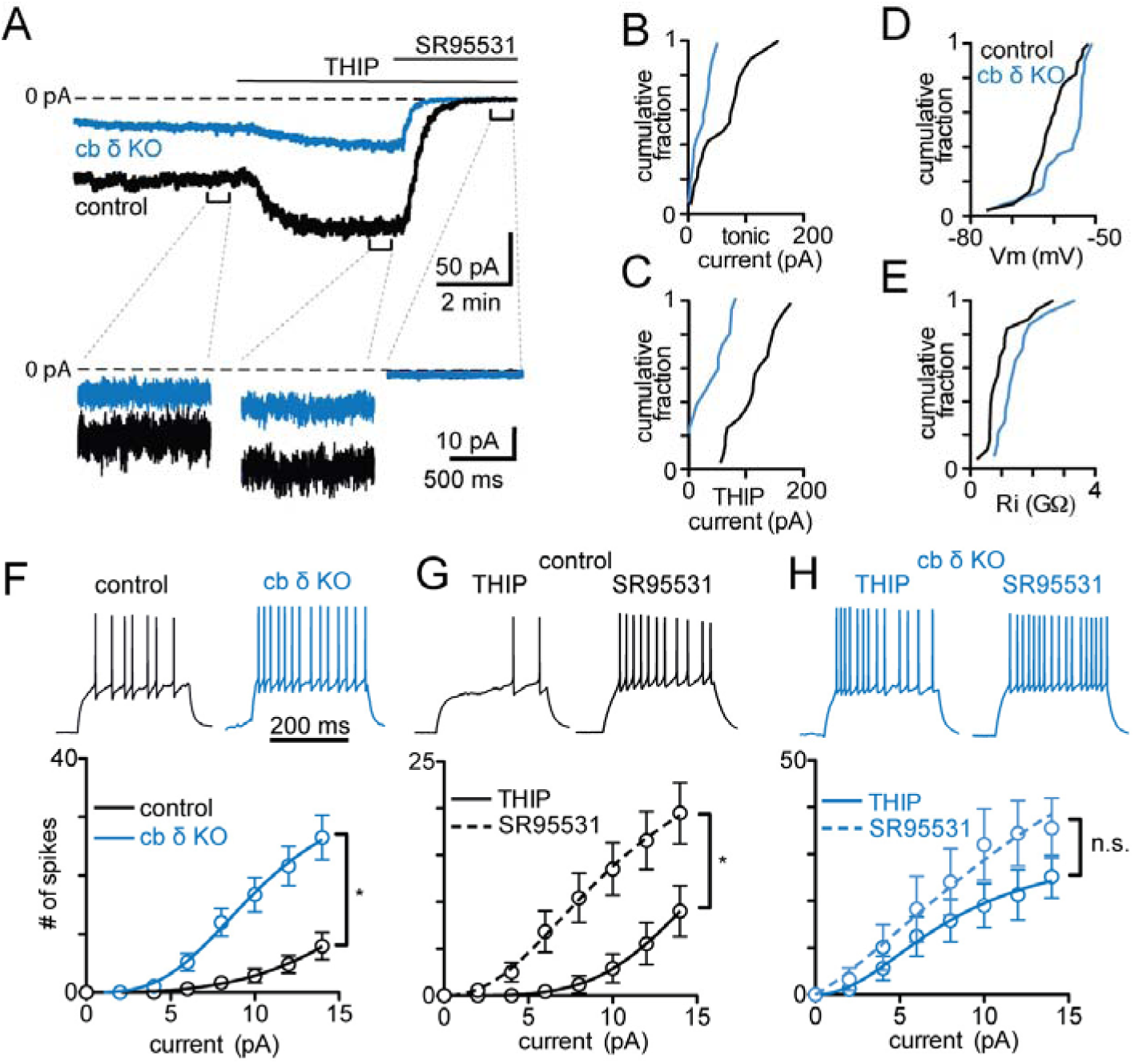
Hyperexcitability and decreased tonic inhibition in granule cells lacking the GABA receptor δ subunit. A) Top, example traces of whole-cell voltage clamp recordings of tonic currents measured during a baseline period, in the presence of the δGABA_A_ subunit containing-preferring GABA_A_R agonist THIP, and in the presence of the non-selective GABA_A_R antagonist SR95531 for a control (black) and a cb δ KO granule cell (blue). Traces below, current traces on an expanded time scale show varying in noise levels depending on the recording condition. SR95531 strongly reduces noise levels compared to control and THIP. B) Cumulative histograms of initial tonic currents in control (black line) and cb δ KO GCs (blue line). C) Cumulative histograms in the presence of THIP. D) Cumulative histograms of membrane potential (V_m_). E) Cumulative histogram of input resistance (R_i_). F) Top, example traces of whole cell current clamp recordings in response to a 10 pA depolarizing current injection in cb δ KO GCs (blue) and in control GCs (black). Bottom, summary data showing the input-output relationships for a range of depolarizing current steps and the resulting number of action potentials. G) As in F but for control animals in the presence of THIP and with GABA_A_Rs blocked. H) As in G but for cb δ KO mice.

Tonic currents regulate GC excitability, but it is not clear whether deletion of δGABA_A_ leads to hyperexcitability, or if compensatory mechanisms counteract the loss of tonic inhibition (Brickley et al., 2001). We assessed GC excitability and found that depolarizing current steps evoked more action potentials in cb δ KO GCs than in control GCs (Figure 2 F, top). This was true for a range of amplitudes (Figure 2F, bottom, control: n=14, cb δ KO: n=14, p<0.0001, 2-way ANOVA), indicating that cb δ KO GCs are more excitable than control GCs. The observations that THIP decreased excitability (Figure 2 G, top left black trace), while inhibiting GABA_A_Rs increased excitability of control GCs (Figure 2 G, top right black trace; bottom: summary input-output curve, control THIP n=19, control SR95531: n=17, p<0.0004, 2-way ANOVA; Figure 2 G), which confirmed the importance of tonic GABA_A_ currents in regulating GC excitability. Conversely, neither activating nor inhibiting GABA_A_Rs had a significant effect on excitability in cb δ KO GCs (Figure 2 H, THIP: n=17, SR95531: n=15, p>0.9, 2-way ANOVA). These experiments indicate that GCs of cb δ KO mice are hyperexcitable because of reduced tonic inhibition.

The GC hyperexcitability evident in cb δ KO mice was surprising because eliminating a GABA_A_ receptor subunit that forms heteromers with δGABA_A_, α6GABA_A_, did not result in hyperexcitability. Rather, despite a strong attenuation of tonic GABA_A_ currents in GCs compensatory mechanisms maintained normal GC excitability (Brickley et al., 2001). Thus, cb δ KO mice provide for the first time the opportunity to assess the behavioral consequences of hyperexcitability of the input layer of the cerebellar cortex.

Cerebellar dysfunction is often associated with ataxia and abnormal fine motor coordination (Manto et al., 2012). We therefore hypothesized that a considerable increase in excitability of the cerebellar input layer in cb δ KO mice could lead to motor dysfunction, including abnormal gait and deficits in motor learning. However, gross motor deficits have not been described in global δ KO mice (Wiltgen et al., 2005), and we found that the same was true for cb δ KO mice (Figure S2 A, B). To examine whether δGABA_A_ deletion results in subtle effects on motor function we analyzed the gait of cb δ KO mice in detail. A high speed, high resolution imaging system allowed us to record and track walking mice and extract the positions of the four paws, the base and tip of the tail and the nose over time (representative frames of recorded movie in Figure 3 A). A velocity plot of different mouse body parts illustrates that control and cb δ KO mice move similarly. The nose and tail advance at a roughly constant velocity, while the hind right/ front left and the front right/ high left paws move together and are either in touch with the floor or moving forward rapidly (Figure 3 B). There was no difference in any parameter describing gait, including cadence, stride length, paw width and stance (Figure 3 C-F, top). As expected, these parameters depend on velocity (Figure 3 C-F bottom), but the velocity dependence in any of these parameters was the same in control and cb δ KO animals. We also analyzed properties of the tail movement and found that tail oscillations that accompany walking, and tail elevation were unaltered in cb δ KO mice (Figure 3 G, H). Finally, we analyzed the fraction of the cycle in which there are a given number of paws on the floor. At low velocities, it is not uncommon that 4 paws are in contact with the floor. As the velocity increases, progressively fewer paws contact the floor, to the point that at very high velocities a fraction of the time there are no paws in contact with the floor (Figure 3I, J). Previous studies have shown that for a given velocity, mice with gait abnormalities tend to have more feet in contact with the floor (Machado et al., 2015). However, we find that there was no difference between control and cb δ KO mice.

**Figure 3:**
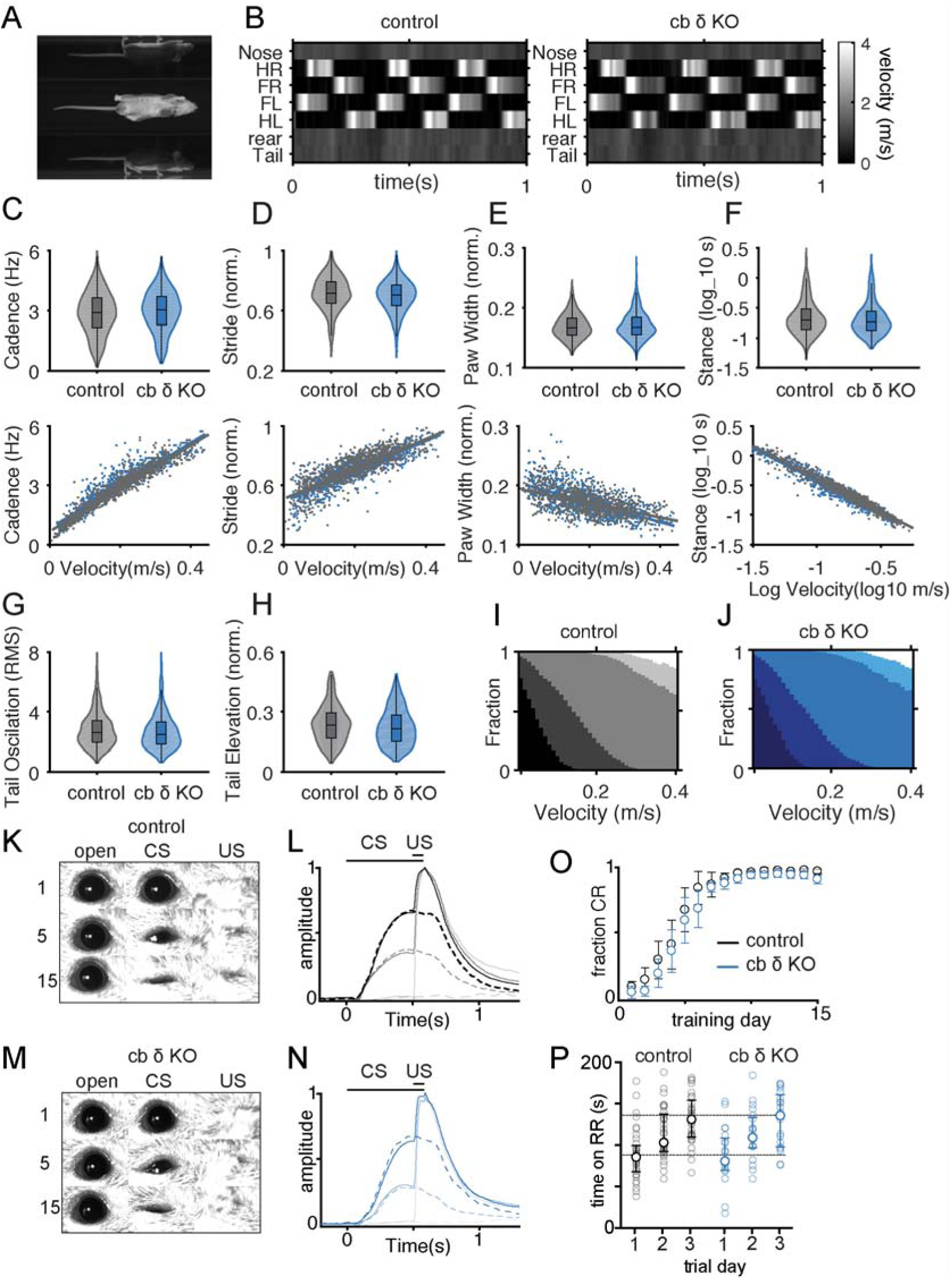
Normal motor function and cerebellar learning in cb δ KO mice. A) sample frame of a walking mouse viewed from the left, right and bottom. B) velocity of different body parts over time between control and cb δ KO. C-F) Top, box and whisker plots overlaid on the density plots for cadence (C), stride (D), paw width (E), stance (F), tail oscillation (G) and tail elevation (H) measured stride-to-stride and pooled across all subjects, with control in grey and cb δ KO in blue. Bottom, scatter plots and linear regression of cadence, stride, paw width and stance against velocity. I,J) fraction of time during a single gait cycle for a given velocity that had four (darkest) to zero (white) supports on the floor. K) Representative images of eye-blink responses in a control animal sampled at the beginning (day 1, top), in the middle (day 5, middle) and at the end of training (day 15, bottom). The baseline, conditioned response (CR) and air puff response (UR) are presented in sequence column-wise. Initially (day 1), the eye remains fully open during the presentation of the CS, and closes fully only with presentation of the US. With continued learning (day 5), the mouse begins to close his eye in response to the CS, until near-full closure at the end of the training period (day 15). The US always elicits a complete closure of the eye. L) The average normalized eye closure from an early (day 1, light grey), middle (day 5, grey) and end of training (day 15, dark grey). t=0 marks the presentation of the CS, and 0.5 s the presentation of the US. Solid lines represent paired presentation of CS and US at different learning stages, and dashed lines represent CS presentation only. M-N) analogous to K, L but in cb δ KO animals. O) Fraction of trials with a CR on each training day. Cb δ KO animals (blue circles) learn at a similar rate as controls (white circles) in an eye blink task (control: n=8, cb δ KO n=8). Data as presented as mean and SEM) P) Motor learning as assayed in a 3-day accelerating rotarod paradigm is similar in control (black) and cb δ KO animals (blue; control: n=36 animals, cb δ KO: 20 animals, p>0.9, Mann-Whitney test). Grey circles represent individual animals.

The cerebellum has a well-established role in motor learning. We therefore tested whether motor learning is disrupted in cb δ KO mice. Eyeblink conditioning is an established learning paradigm that requires the cerebellum (Thompson, 1986). A conditioning stimulus (CS, a weak illumination) that does not initially cause the eye to close, is paired with an unconditioned stimulus (US, air puff to the eye) that causes the eye to close. Delay conditioning experiments were performed in mice running on a rotating platform as the CS and US were presented (Albergaria et al., 2018). The CS occurs for 500 ms, and for the last 50 ms it is paired with the US. On the first day of CS/US pairing, the CS alone does not cause eyelid closure, by the fifth day the CS alone causes the eyelid to partially close in a fraction of trials, and by the fifteenth day of pairing, mice close their eyes in response to the CS alone in most trails. As shown in example trials, cb δ KO mice learn similarly to control animals (Figure 3 control: K, L; cb δ KO: M, N). All control mice and cb δ KO mice learned to respond to the CS, and control and cb δ KO mice both learned at approximately the same rate (Figure 3 O, control: n=8, cb δ KO: n=8, see table 1). Average responses to the CS alone and to paired CS + US stimulation were very similar for control and cb δ KO mice on days 5 and 15 (Figure 3 L, N). Since cerebellar deficits can affect motor learning (Kloth et al., 2015; Tsai, 2016) we also performed a 3-day accelerating rotarod paradigm. We found that there was no difference between control mice and cb δ KO mice in initial performance on the rotarod (control, 90 ± 5 s on rotarod, n=36, cb δ KO, 89 ± 8 s on rotarod, n=20, p>0.9, Mann-Whitney test; Figure 3 P), and this was the case for both females and males (Figure S2 C). Improvement of motor performance in control and cb δ KO animals was similar by the end of training (Figure 3 P, control: 136 ± 6 s on rotarod, n=36; cb δ KO 133 ± 7 s on rotarod, n=20, p>0.9, Mann-Whitney test). Thus, remarkably, despite hyperexcitability of the cerebellar input layer in cb δ KO mice, we did not detect any gait abnormalities or defects in motor learning.

It has been shown previously that some cerebellar deficits cause neuropsychiatric symptoms without producing major motor deficits. We therefore used Motion Sequencing, MoSeq, (Wiltschko et al., 2015) to test whether cb δ KO mice show behavioral abnormalities that are difficult to detect with alternative approaches. MoSeq posits that spontaneous behavior consist of discrete and stereotyped three-dimensional motifs of behavior called “syllables,” and it identifies and quantifies these syllables using unsupervised machine learning. MoSeq can detect subtle behavioral differences caused by genetic mutations in mouse models of disease, by sensory cues or changes in the environment, and in response to optogenetic manipulation (Markowitz et al., 2018; Pisanello et al., 2017; Wiltschko et al., 2015).

Single mice were observed in a circular behavioral arena with a depth camera (Figure 4 A, left). Syllables were extracted offline, as shown for three sample syllables (Figure 4 A, right). MoSeq identified over 90 distinct syllables (Figure S6). A comparison of the 30 most common syllables identified a subset of syllables whose usage differed between control and cb δ KO animals (Figure 4 B). Although many syllables were indistinguishable between control and cb δ KO animals, pausing and stretching syllables were used less in cb δ KO animals (Figure 4 B, C). Conversely, the usages of syllables corresponding to swift movements, such as a dart or a rapid down turn, were elevated in cb δ KO animals (Figure 4 B, E). A subset of these syllables also showed increased durations, although most syllables had similar durations in control and cb δ KO mice, regardless of whether they were differentially used or not (Figure S3 A, B). MoSeq also identified less common behaviors that were more frequently used in cb δ KO animals, including long-lasting bouts of grooming and high jumps on the walls of the bucket, presumably to escape the arena (Figure 4 F). Thus, although the syllabic repertoire of control and cb δ KO animals is similar, syllable usage patterns differ between the genotypes. Decreased pauses, increased rapid movements, hyperlocomotion, increased grooming and more escape attempts in cb δ KO mice are consistent with an increase in anxiety-related behaviors, reminiscent of complex behaviors displayed after stress (Füzesi et al., 2016), and a general inability to calm down after being placed in the novel environment of the behavior arena. This observation is particularly interesting because the δGABA_A_ subunit has been implicated in these behaviors (Liu et al., 2017; Maguire et al., 2005; Shen et al., 2007; Zhang et al., 2017b), and δGABA_A_ modulators have been proposed to reduce anxiety in humans (Ströhle et al., 2002), but the possibility that δGABA_A_ in the cerebellum could contribute to anxiety-like behaviors has not been previously demonstrated.

**Figure 4:**
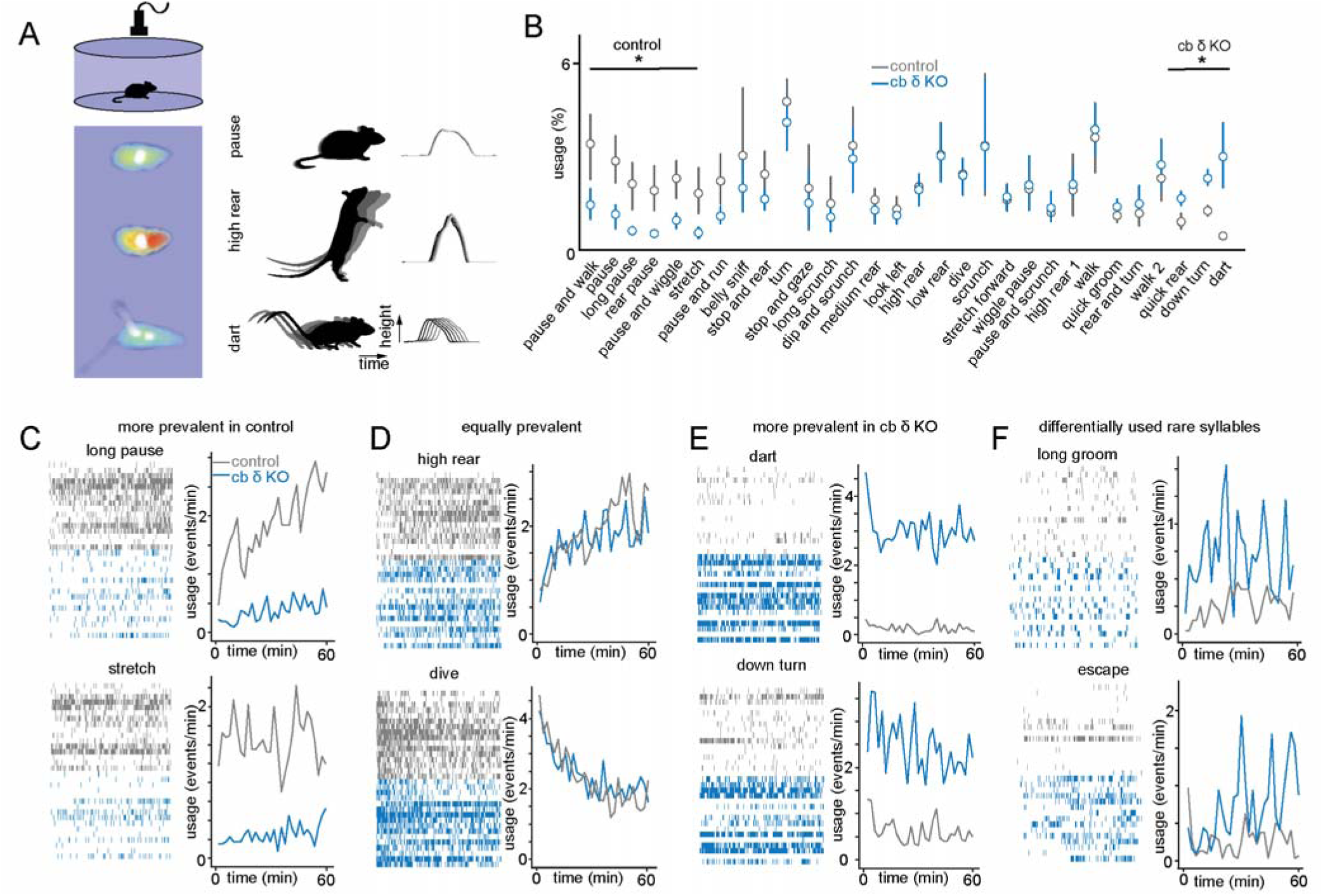
Unsupervised analysis reveals behavioral abnormalities in cb δ KO animals. A) 3D imaging of mice in the open field with a depth camera, followed by MoSeq-based segmentation of the behavior data, reveals behavioral syllables (see methods); example behavioral syllables and the associated inferred positions of the spine in time and space are in (middle and right). B) A usage plot for the 30 most frequently-used syllables. A subset of behavioral syllables is differentially expressed in control and cb δ KO animals. C) Example syllables performed more frequently by control animals. Left, raster plots of syllable occurrence during observation period, right, syllable usage over time. The observed behavior associated with each syllable is indicated. D) Like (C) but depicting common example syllables that are equally prevalent in both control and cb KO. E) Like (C) but depicting common example syllables performed more frequently by cb KO animals. F) Like (C) but depicting rarely used syllables that are more frequently used by cb KO animals.

Anxiety-related behaviors in rodents are elicited by aversive and unpredictable external stimuli that can occur acutely and for short durations, or chronically over extended periods of time. The behavioral expressions are complex, but can include vigilance and changes in locomotion, as we observed with MoSeq. To further examine the effect of δGABA_A_ deletion in the cerebellum on anxiety-related behaviors, we conducted a light/dark test that was previously used to detect differences in anxiety levels in mouse models of disease and in mice treated with anxiolytic drugs (Crawley, 1985; Lezak et al., 2017). This test tracks mice in an arena divided into two compartments, one dimly lit and the other brightly lit (Figure 5 A). It examines whether a mouse overcomes the potential danger of entering a brightly lit space to explore a novel environment, or prefers to remain in a dark space that is perceived as safer. The assumption is that the more anxious the mouse, the less time it spends in the bright compartment. Cb δ KO mice spent less time than control mice in the bright compartment, as shown in median heat maps (Figure 5 B), and in the summaries of individual animals tested (Figure 5 C). Control animals spent 34 ± 3 % (n=32) of the time in the light compartment, compared to 18 ± 3% (n=27) for cb δ KO mice (p<0.0008, Mann-Whitney test). Both males and females exhibited similar differences between control and cb δ KO mice (Figure S5). In addition to a difference in time spent in the light chamber, a higher fraction of cb δ KOs exclusively remains in the dark compartment during the observation period compared to control animals (6/27 and 1/32, respectively).

**Figure 5.**
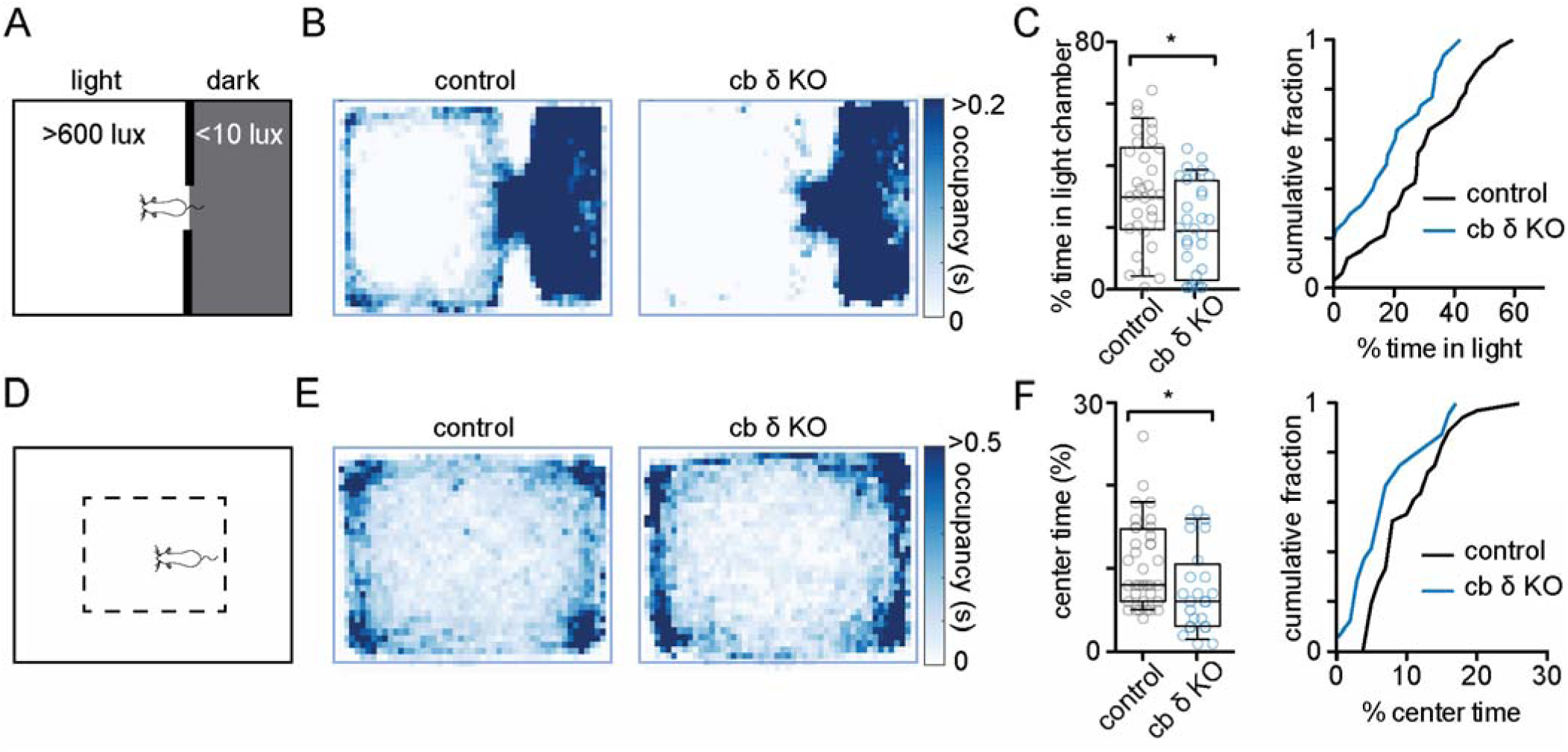
Increased anxiety-like behaviors in cb δ KO animals. A) Schematic of the light/dark chamber. Mice are placed in a behavioral arena divided into a dark compartment (<10 lux) and a lit compartment (>600 lux). A door connects the two chambers, allowing navigation between both compartments. B) Median occupancy plots indicate that cb δ KO animals (*right*) spend less time in the lit compartment than control animals (*left*). C) Summary data of % time spent in the light compartment. Left, boxes denote interquartile range and median, whiskers represent 10-90 percentile. Circles show individual control (n=32, grey circles) and cb δ KO (n=27, blue circles) animals (p<0.009, Mann-Whitney test). Right, cumulative fraction of time spent in the light compartment (control: black line, cb δ KO: blue line) D) Schematic of open field behavior arena. Dashed rectangle denotes center of the area. E) Median occupancy plots of control (left) and cb δ KO animals (right). F) Left, summary data of % time spent in the center of the behavioral arena circles show individual control (n=36, grey circles and cb δ KO (n=24, blue circles) animals (p<0.02, Mann-Whitney test). Right, cumulative probability graph of center time.

We also observed mice in an open field paradigm. Decreased time spent in the center region of the enclosure has been interpreted as elevated levels of anxiety-like behavior (Figure 5 D). Median occupation heat maps (Figure 5 E) and summaries of individual animals (Figure 5 F) show that cb δ KO mice spent less time than cb δ KOs in the center of the behavior arena. On average, cb δ KOs spent 7.3 ± 1 % in the center compared to 10.3 ± 1 % (n=36) for control mice p>0.02, Mann-Whitney test). Taken together, the light/dark test and the open field test suggest that cb δ KO mice display increased anxiety-like behavior compared to control mice.

Taken together, several lines of evidence raised the possibility that δGABA_A_ function in the input layer of the cerebellum could also influence social behaviors. First, global decreases in δGABA_A_ function have been associated with increased fear-related behavior and sensitivity to stress (Maguire and Mody, 2007; Mody and Maguire, 2011; Wiltgen et al., 2005), that in turn have been linked to a diminished interest in social interactions (Beery and Kaufer, 2015; File and Hyde, 1978). This hypothesis is further supported by the notion that anxiolytics acting on δGABA_A_ have prosocial effects (File, 1980). Our findings indicate that loss of δGABA_A_ in cerebellar GCs contributes to increased anxiety-like behavior that could affect sociability. Second, cerebellar dysfunction can lead to social deficits in humans and in animal models (Badura et al., 2018; Carta et al., 2019; Schmahmann and Sherman, 1998; Tsai, 2016). Third, decreased tonic inhibition within the cerebellar input layer are evident in several global mouse models of psychiatric disorders (Bruinsma et al., 2015; Kim et al., 2017).

We therefore examined social behaviors in cb δ KO mice. We used a 3-chamber paradigm in which mice were allowed to navigate between a chamber containing a sex-matched juvenile conspecific (social stimulus, S), an empty middle chamber, and a chamber containing a novel object (O). In the absence of any stimuli, animals did not show a preference for either side of the chamber (Figure S6 A-C). However, median occupation heat maps indicate that control mice prefer investigating the social stimulus to the object, while cb δ KO mice show no preference for either stimulus (Figure 6 A, control: left; cb δ KO: right). Summary data of individual animals is shown in Figure 6 B-C (control, 32 ± 1 % in O, 50 ± 1% in S, n=54, p<0.0001; cb δ KO, 37 ± 2 % in O, 44 ± 2 in S, n=29, p>0.1, Mann-Whitney test). Control and cb δ KO animals did not prefer either chamber in the absence of stimuli (Figure S6 A-B). Control animals also investigated the conspecific for a longer duration than cb δ KO mice (Figure S6 C, control 133 ± 9 s, n=54 cb δ KO: 108 ± 9 s, n=29, p<0.04, Mann-Whitney test). But the number of entries to the object or social chambers were similar (Figure S6 D), suggesting that control and cb δ KO did not avoid either chamber. We tested whether the lack of social interest exhibited by cb δ KO mice was due to an olfactory deficit by assessing the response to several non-social (water, coconut, raspberry, banana) and social (male and female urine) odors. Both control and cb δ KO mice explored the odors for similar durations, with the exception of coconut and male urine, in which cb δ KOs showed a slightly increased interest compared to control (Figure S6 E). These experiments indicate that social deficits in cb δ KO mice are not a consequence of an inability to detect social odors.

**Figure 6:**
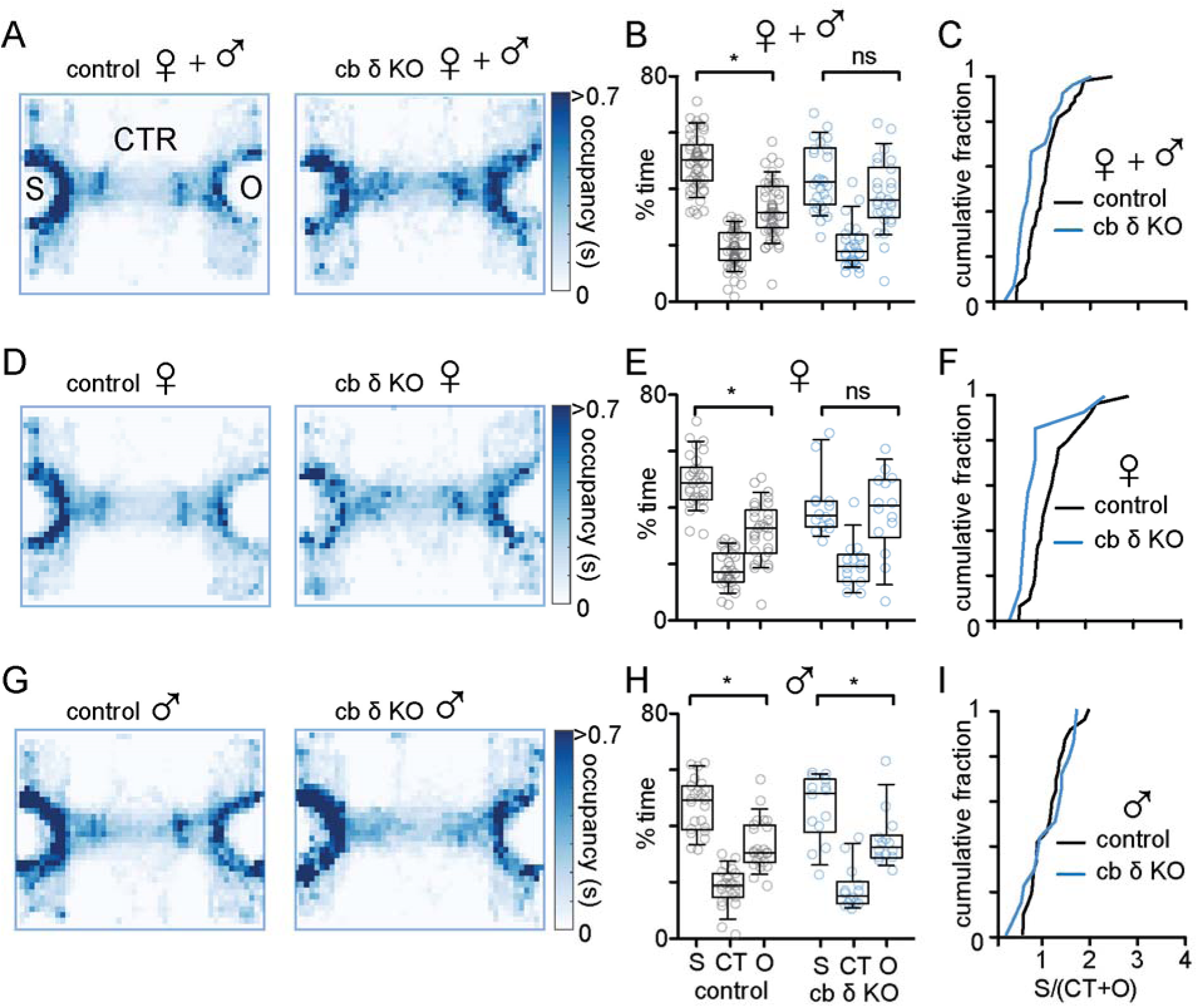
Sexually dimorphic effect of cerebellar-specific δGABA_A_ deletion on social behavior. A) Median occupancy plots indicate a clear preference of control (*left*) animals for the social stimulus over the object, while cb δ KO (*right*) animals show no preference for either stimulus. B) Summary data for % time spent in the object, center and social compartment of the behavior chamber during the observation period. On average, control animals display a strong preference for the social stimulus (grey circles, p<0.0001), while cb δ KO animals show no preference (blue circles) C) Cumulative probability plot of the ratio of time spent with the social stimulus and the sum of the time spent in the center (CTR) and object chambers (O); sociability ratio S/(O+C), control: black trace; cb δ KO, blue trace) D) – F) same as A) – D) but females only. D) median occupancy plots of control (*left*) and cb δ KO females (*right*). Control females show a strong preference for the social stimulus (*left*) while cb δ KO (*right*) females do not show a preference for either stimulus. E) Summary data of % time spent in social, center and object compartment. F) Cumulative probability plot of sociability ratios in females. G) - I) same as A) – D) but males only. G) Median occupancy plots of control (left) and cb δ KO males (right). Both control and cb δ KO males prefer the social stimulus over the object stimulus. H) Summary data of % time spent in social, center and object compartment. I) Cumulative probability plot of the sociability ratios in males.

Separate analysis of females and males revealed that in cb δ KO mice social deficits were pronounced in females and absent in males. Median occupancy heat maps for females show that control mice strongly preferred the social chamber over the object chamber, but cb δ KO mice had no preference (Figure 6 D). Figure 6 E-F show the summary data of individual experiments (control, 32 ± 2 % in O, 50 ± 2 % in S, n=29, p<0.0001; cb δ KO, 39 ± 4 % in O, 41 ± 3 in S, n=15, p>0.8). In contrast, for males, control and cb δ KO mice showed a similar preference for the social chamber (Figure 6 G-I, control, 33 ± 2 % in O, 48 ± 2 % in S, n=25, p<0.002; cb δ KO, 34 ± 3 % in O, 48 ± 3 in S, n=14, p<0.04, Mann-Whitney test). No significant differences between males and females were observed in responses to odors (Figure S6 F). Thus, deletion of cerebellar δGABA_A_ subunits in GCs leads to social deficits in females but not in males.

The neurosteroid sensitivity of δ subunit-containing GABA_A_Rs could contribute to the sex dependence of social deficits in cb δ KO mice. This raises the possibility that δGABA_A_-containing receptors in the cerebellum might also contribute to other behaviors that involve the δGABA_A_ subunit and steroid-dependent modulation in females, such as postpartum-related changes in in anxiety and maternal care (Maguire and Mody, 2008). Little is known about which δGABA_A_-expressing brain regions mediate these behaviors, but it had been assumed that the cerebellum is not involved. Given the high expression levels of δGABA_A_ in the cerebellum, social deficits in cb δ KO females specifically, and increased anxiety in cb δ animals that could contribute to poor maternal care, we tested parental behavior in cb δ KO females. We also included heterozygous cb δ females (cb δ HET) because parental behavior is compromised in heterozygous global δ KOs (Maguire and Mody, 2008). Newborn pups emit olfactory signals and ultrasonic vocalizations that can stimulate spontaneous parental behavior in virgin females including licking and crouching over pups (Calamandrei and Keverne, 1994; Gandelman, 1973; Noirot, 1969; Thomas and Palmiter, 1997). We thus first tested whether δGABA_A_ deletion in the cerebellum affects spontaneous parenting in virgin females. We developed a test in which female virgins were allowed to interact with a nest of newborn (P1-P3) pups and inanimate pup-sized objects to control for novelty (Figure 7 A). Median occupation heat maps (Figure 7 B) and summaries of individual experiments (Figure 7 C-F) show that while control females interact almost exclusively with pups, cb δ HET and KO females were less interested in pups and also increasingly interacted with objects. Upon habituation with pups, female virgins are known to retrieve displaced pups to the nest (Noirot, 1969), Figure 7 H). We found that the control virgins retrieved at higher rates than cb δ KO and cb δ HET virgins (Figure 7 G, control 43 %, cb δ HET 29 %, cb δ KO 5%, p<0.009, Chi-square test). These results suggest that virgin cb δ KOs and cb δ HET females are less likely to express spontaneous parental-like behavior towards pups.

**Figure 7:**
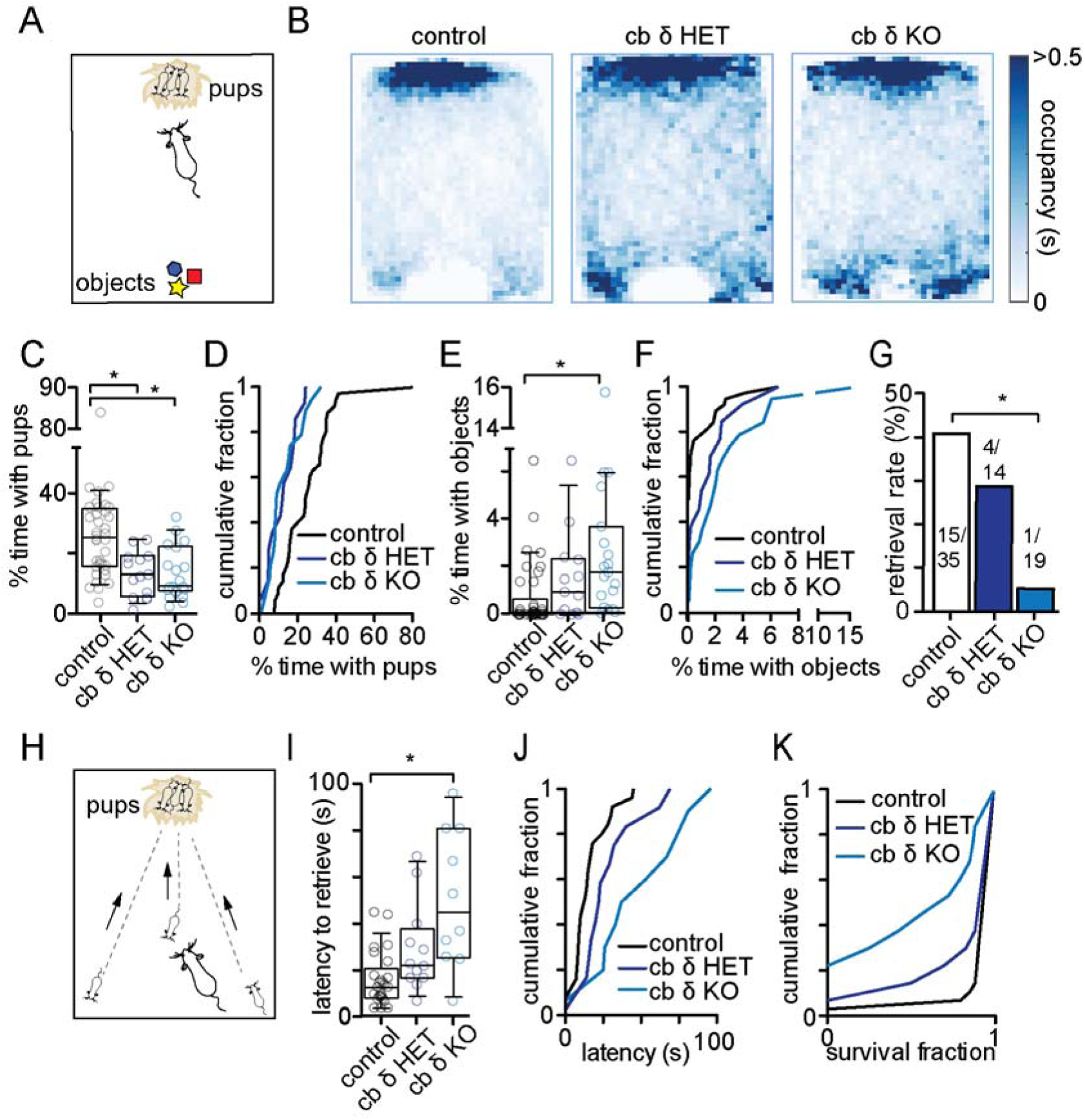
Cb δ KO females display altered maternal behavior. A) Schematic of behavioral paradigm. A virgin female is placed in the behavioral arena that contains a nest with three pups (top) and three pup-sized novel objects (bottom). B) Median occupancy plots of control, cb δ HET and cb δ KO females. Control females preferentially remain within close proximity of the nest while cb δ HET and cb δ KO mice disperse around the nest and explore objects. C) Summary plot of time spent with pups during the observation period. Circles show individual mice tested (control: grey, n=37, cb δ KO het: dark blue, n=12, cb δ KO: blue, n=19, Kruskal-Wallis test with Dunn’s post-test, p<0.003). D) Cumulative probability plot of time spent with pups. KS test, p<0.05 E) Summary plot of time spent with pup-sized objects. cb δ KO mice show a greater interest in objects than controls. (control: grey, n=37, cb δ HET: dark blue, n=12, cb δ KO: blue, n=19, Kruskal-Wallis test with Dunn’s post-test, p<0.006) F) Cumulative probability plot of time spent with objects. KS test, p<0.05 G) Retrieval rate in control, cb δ HET and cb δ KO virgin females (control: n=35, cb δ HET: n=14, cb δ KO: n=19, p<0.009, Chi-Square test) H) Schematic of retrieval paradigm. In the home cage, 3 pups are removed from the nest and placed in the opposite 2 corners and in the center of the cage. I) Summary plot of latency to retrieve the first pup. Circles show individual mice tested (control: grey, n=25, cb δ KO het: dark blue, n=12, cb δ KO: blue, n=10, Kruskal-Wallis test with Dunn’s post-test, p<0.008) J) Cumulative probability plot of retrieval times (KS test p<0.008) K) Cumulative probability histogram of pup survival fraction.

To assess whether cerebellar deletion of δGABA_A_ alters postpartum behavior we first assayed a robust behavior commonly expressed by postpartum mice, the retrieval of displaced newborn pups to the nest (Figure 8 H). We found that cb δ KO dams delayed initiation of retrieval compared to controls (control: 16 ± 2 s, n=25; cb δ HET: 29 ± 5, n=12; cb δ KO: 51 ± 9 s, n=10, p<0.008, Kruskal-Wallis test, Figure 8 I-J). cb δ KO dams also took longer to complete retrieval of three displaced pups (Figure S7 A, B). In addition, similar to previous reports of global δ KO females (Maguire and Mody, 2008), we observed increased cannibalization of newborn pups in cb δ KO females (image pups of representative litters in Figure S7 E), resulting in a reduced survival fraction of pups (Figure 7 K). We did not observe any differences in fertility or litter sizes (Figure S7 C-D), indicating that reproduction was not affected in cb δ KOs. Counter to global δ KO dams (Maguire and Mody, 2008), we found no difference in the Porsolt forced swim test (Figure S7 G), or failure to build nests (data not shown) in cb δ KO females. Taken together, these results suggest that a subset of the abnormal maternal behaviors in global δ KOs were also observed in cb δ KOs and cb δ HET KOs.

## Discussion

Here, we find that the cerebellum, traditionally viewed as having largely motor functions, regulates behaviors that are relevant to many psychiatric and neurodevelopmental disorders. We show that specific deletion of δGABA_A_ from cerebellar GCs attenuates tonic inhibition and increases excitability in the input layer of the cerebellar cortex, which in turn produces diverse and unexpected non-motor deficits without disturbing motor behavior. Our study establishes that the cerebellar input layer is a critical processing stage for controlling anxiety-related behaviors, social behavior in females, and maternal care.

### Hyperexcitability and lack of dynamic regulation of tonic inhibition in GCs of cb δ KO mice

The pronounced hyperexcitability (Figure 2) that accompanies δGABA_A_ elimination from GCs is striking given that eliminating α6GABA_A_ fails to alter GC excitability due to upregulation of 2-pore K+ channel upregulation that offsets the loss of tonic inhibition (Brickley et al., 2001). It is not known why compensatory K^+^-channel expression in GCs is prominent in α6 KO mice, but not in cb δ KOs. In addition, the GABA_A_ receptors of GC axons, the parallel fibers (PFs), are thought to control axonal excitability and transmitter release (Pugh and Jahr, 2011). However, global δGABA_A_ deletion was reported not to change PF excitability (Dellal et al., 2012). It is therefore unlikely that PF-mediated excitation of Purkinje cells (PCs) is changed in cb δ KO mice.

Another important consequence of eliminating δGABA_A_ from GCs is their diminished ability to respond to contextual cues, provided my modulatory neurotransmitters, neuropeptides and neurosteroids. Neuromodulators can influence ambient GABA levels by regulating firing of spontaneously active Golgi cells and PCs that control tonic inhibition (Duguid et al., 2012; Farrant and Nusser, 2005; Guo et al., 2016; Mitchell and Silver, 2003). Due to their high GABA affinity, δ/α6GABA_A_ containing receptors can faithfully detect changes in ambient GABA levels. Golgi cell firing has been shown to depend on synaptic input and neuromodulators, such as serotonin and acetylcholine (Fleming and Hull, 2019; Hull et al., 2013), that allow fine-tuning of extrasynaptic GABA levels, tonic inhibition and GC excitability. In addition, neurosteroids modulate δGABA_A_-containing receptors directly (Stell et al., 2003; Vicini et al., 2002) and can regulate GC excitability during diverse physiological states, such as during the estrous cycle (Maguire et al., 2005; Wu et al., 2013), postpartum and during stress (Camille Melón and Maguire, 2016; Maguire and Mody, 2008; Maguire et al., 2009). For these reasons, the ability of GCs to dynamically adjust their excitability in an activity and context-dependent manner is compromised in cb δ KOs.

### Motor function is normal in cb δ KO mice

Given the cerebellum’s heavily studied role in locomotion, we were surprised that no motor phenotype in cb δ KO mice was detectable (Figure 3). There is precedence for motor performance being relatively insensitive to manipulations of the cerebellar input layer. Motor deficits were also mild or absent in several mouse models that describe deficits in tonic GC inhibition (Bruinsma et al., 2015; Wiltgen et al., 2005). However, it is not known whether GCs are hyperexcitable in these mice, or if compensatory mechanisms are prominent, as in α6 KO mice. It has also been shown that strongly attenuating the strength of GC synapses also did not affect basic locomotion (Galliano et al., 2013) but degraded several forms of cerebellar learning. These observations suggest that basic motor function is remarkably insensitive to manipulations of the input layer, but that a decrease in the influence of the GC layer may be more detrimental to motor learning than hyperactivation.

Several explanations could account for the lack of motor deficits in cb δ KO mice. First, it is possible that cerebellar circuitry, in which GCs both excite PCs directly and disynaptically inhibit PCs by activating molecular layer interneurons, can overcome GC hyperexcitability by increasing inhibition of PCs by MLIs. Second, perhaps circuit-level compensation can overcome the loss of tonic inhibition in cb δ KO mice. Specifically, even though upregulation of K-channels in GCs is not prominent in cb δ KO mice, it is possible that mossy fiber to GC synapses or GC synapses onto their targets become weaker, or that circuitry is altered within the cerebellar cortex or in downstream brain regions.

It is particularly interesting that motor performance was insensitive to decreased tonic inhibition, whereas non-motor behaviors were so strongly affected. Changes in conditioned eyeblink have been observed in numerous autism mouse models (Kloth et al., 2015) and in patients suffering from neurodevelopmental disorders (Reeb-Sutherland and Fox, 2015), suggesting that non-motor and motor deficits often go hand in hand. In mice in which cerebellar PCs are selectively perturbed (Pcp2-Cre x flox/+ Tsc1) (Tsai et al., 2012; 2018), social deficits were observed, locomotion appeared normal, but subtle deficits in conditioned eyeblink were apparent. Here, we find increased anxiety-like behavior, altered social behaviors and maternal care in cb δ KO mice, but gait and motor learning were unaffected. Each of these behaviors are thought to involve different regions of the cerebellar cortex (Krienen and Buckner, 2009; Stoodley and Schmahmann, 2018). It is possible that these regions perform somewhat different computations, or that parts of the cerebellar cortex and downstream brain regions are specialized in ways that make them more or less sensitive to GC hyperexcitability.

### Increased anxiety-like behavior in cb δ KO mice

Unbiased behavioral testing with MoSeq is well-suited to characterizing behavioral alterations arising from manipulations of the cerebellum, which controls both motor and non-motor behaviors. By generating a detailed profile of spontaneous behavior with MoSeq, we found that cb δ KO mice were more restless, displayed increased darting and escape attempts, and decreases in calm, restful behaviors, like pausing and stretching (Figure 4). These findings are consistent with a hyperlocomotion phenotype, and similar hyperactivity was reported in mice treated with stimulants (Wiltschko et al. in revision), in mouse models of ASD and ADHD (Angelakos et al., 2017; Dalla Vecchia et al., 2019; Schmeisser et al., 2012), and in response to stress (Füzesi et al., 2016; Strekalova et al., 2005). Our subsequent experiments using the light/dark chamber and open field assays were consistent with cb δ KO mice showing both increased levels of anxiety-like behavior and hyperactivity. Although freezing and hypolocomotion are commonly associated with an anxious behavioral state in mice, hyperactivity in conjunction with anxiety have been reported in some instances, and anxiety is a common comorbidity of ADHD and ASD (Kazdoba et al., 2016).

The elevated anxiety-like behavior in cb δ KO mice is particularly interesting in light of the many clinical reports implicating the cerebellum in anxiety, and considerable evidence suggesting that δGABA_A_ plays an important role in anxiety and fear-related behaviors. The cerebellum is associated with phobia, generalized anxiety disorder and PTSD (Caulfield et al., 2016; Moreno-Rius, 2018). Much less is known about the cerebellum and anxiety in animal models. *Lurcher*, a PC degeneration mouse model of ataxia, shows an abnormal fear response (Hilber et al., 2004; Lorivel et al., 2014). However, although this model is a gain-of-function mutation of GluRδ2 that is expressed at particularly high levels in PCs, it is also found elsewhere in the brain. Several lines of evidence suggest that δGABA_A_Rs also play an important role in anxiety. Endogenous and pharmacological compounds that preferentially act on δGABA_A_Rs affect anxiety (Eser et al., 2006). Fluctuating levels of neurosteroids and δGABA_A_ expression are thought to contribute to anxiety and mood swings during the diestrus phase of the ovarian cycle (Maguire et al., 2005; Smith et al., 2006) and puberty (Smith, 2013). Ethanol is thought to produce anxiolytic effects by increasing tonic inhibition synergistically by both directly targeting extrasynaptic δGABA_A_Rs and by enhancing Golgi cell firing, which raises GABA levels in the cerebellar input layer (Richardson and Rossi, 2017). Previous studies investigating δGABA_A_Rs and anxiety focused on brain regions classically associated with fear, such as the amygdala, hypothalamus and hippocampus (Lee et al., 2014; Liu et al., 2017; Maguire et al., 2005). Our study establishes the importance of δGABA_A_Rs in the input layer of the cerebellum in regulating anxiety.

Elevated levels of anxiety are also linked to many neurological disorders such as ASD, ADHD, and postpartum depression. Compelling clinical evidence suggests that the cerebellum plays an important role in many psychiatric and neurodevelopmental disorders, such as schizophrenia, ADHD and ASD (Sathyanesan et al., 2019; Stoodley, 2016), presumably because of extensive connections between the cerebellum and brain regions involved in executive function, cognition and emotional control (D’Mello and Stoodley, 2015; Strick et al., 2009; Wang et al., 2014). Our results indicate that perturbation of the input layer of the cerebellum can elevate anxiety and stress, and this could contribute to all of these cerebellar-dependent behaviors. In addition, elevated anxiety-like behavior in cb δ KO mice could contribute to deficits in maternal behaviors and postpartum depression.

### Sex-specific effects on sociability in cb δ KO mice

The changes in social behavior of cb δ KO mice that were apparent in the 3-chamber paradigm (Figure 6) could arise in many ways. In mice, social behavior relies mainly on olfactory and auditory cues emitted by interacting conspecifics, but it is unlikely that social deficits in cb δ KO mice reflect differences in primary sensory processing. We found no difference in odor recognition (Figure S6 G), and hearing is normal in global δ KO mice (Maison et al., 2006). The social deficits observed in our cerebellar-specific manipulation are consistent with the considerable evidence for the cerebellum’s involvement in social behavior, e.g. in the context of ASD and cerebellar cognitive affective syndrome (Schmahmann, 2019; Tsai, 2016; Wang et al., 2014). What sets our findings apart from previous cerebellar-specific manipulations is that the social deficits were observed exclusively in female mice. In addition, rather than targeting the output of the cerebellar cortex (PCs), we have manipulated the input layer of the cerebellar cortex (GCs). The sex specificity is particularly interesting because several psychiatric disorders, such as depression and schizophrenia, show sex bias in humans (Werling and Geschwind, 2013). Our findings suggest that treatments that target δGABA_A_Rs, such as the recently introduced post-partum depression drug brexanolone (Zulresso^TM^, Sage Therapeutics) might provide an effective treatment of deficits associated with neurodevelopmental disorders in females. Clinical trials of drugs targeting δGABA_A_Rs in Angelman syndrome, a non-dimorphic form of autism, are already underway (STARS, Bird, Ochoa-Lubinoff, Tam et al., 2019 American Academy of Neurology Annual Meeting; May 4-10, 2019; Philadelphia, PA)

Their neurosteroid sensitivity makes δGABA_A_Rs well suited to mediating sex-specific differences in physiology and behavior. Sexually dimorphic behavioral deficits could arise from different baseline levels of gonadal hormones, neurosteroids and synthesis enzymes in females. These could modulate tonic GABA_A_ currents and make females more vulnerable to δGABA_A_ deletion (Schüle et al., 2014; Ströhle et al., 2002; Yoshizawa et al., 2017). The observations that baseline tonic currents are larger in females but THIP-evoked currents are of equal amplitude are consistent with this hypothesis (Figure S1 G, H). Another possibility is the well-established developmental influence of gonadal hormones on social behavior (Bell, 2018). The lack of δGABA_A_ in GCs during development could decrease sensitivity to gonadal hormones, and this could affect males and females differently. In addition, it is likely that elevated anxiety-like behavior in cb δ KO mice decreases interest in social interaction (File and Hyde, 1978). Although anxiety-like behavior was evident in both cb δ KO males and females, the effect on social behaviors could differ. Evidence that stress can result in sexually dimorphic effects has been reported elsewhere (Senst et al., 2016) and could contribute to behavioral differences. Finally, protein deletion can affect gene expression in a sex-dependent manner, as occurs in Angelman syndrome (Koyavski et al., 2019). Sex differences in functional compensation have been observed in the cerebellum, as in an ASD mouse model at the level of the DCN, the first output targets of the cerebellar cortex (Mercer et al., 2016).

### Abnormal maternal behavior in cb δ KO females

There is considerable evidence for the involvement of δGABA_A_Rs and neurosteroid signaling in postpartum depression (Maguire and Mody, 2009). The surge of progesterone metabolites during early pregnancy, followed by their decline before giving birth, regulate δGABA_A_R function and expression levels (Maguire and Mody, 2008). The rapid upregulation of δGABA_A_Rs is thought to curb postpartum depression-like behaviors like infanticide and anxiety, and these behaviors that were prominent in postpartum global δ KO mice dams. However, conditional δ KO mice have not been used previously to examine postpartum depression-like behaviors, and the brain regions regulating postpartum depression have not been identified. The cerebellum was never considered to be a likely candidate because there had never been any suggestion that it might be involved in such behaviors. Instead upregulation of δGABA_A_ expression in dentate GCs (Maguire and Mody, 2008; Maguire et al., 2009) and in hippocampal PV+ interneurons (Ferando and Mody, 2013) have been implicated in the regulation of postpartum depression, although this was never directly tested in conditional δ KO mice.

Unexpectedly, we observed abnormal maternal care in cb δ KO mice, including lack of interest in pups, failure to retrieve and increased cannibalization of pups (Figure 7). Given the strong connection between anxiety and stress in the genesis of postpartum depression (Camille Melón and Maguire, 2016; Payne and Maguire, 2019), the elevated levels of stress and anxiety in cb δ KO mice are likely a major contributing fact to the observed abnormal maternal care in cb δ KO mice. Our results suggest that some of the behavioral consequences on abnormal postpartum behaviors, like pup retrieval and cannibalization, are comparable for global KOs and cb δ KO mice. Further studies are needed to determine if δGABA_A_Rs in other brain regions also contribute to these behaviors.

In summary, we examine for the first time how hyperexcitability of the cerebellar input layer affects behavior. We find that the repertoire of behaviors influenced by the cerebellum is far greater than previously appreciated. Our results provide insight into how cerebellar dysfunction could lead to the behavioral abnormalities that occur in many psychiatric and neurodevelopmental disorders. Thus, we speculate that manipulating excitability of the cerebellar input layer could relieve some symptoms associated with these disorders.

## Key resources table

**Key resources table:**
| Reagent or Resource | Source | Identifier |
| --- | --- | --- |
| **Chemicals, Peptides, and Recombinant Proteins** |  |  |
| NBQX disodium salt | Abcam Cambridge, MA | Cat# ab120046 |
| (R)-CPP | Abcam Cambridge, MA | Cat# ab120159 |
| SR95531 (gabazine) | Abcam Cambridge, MA | Cat# ab120042 |
| Strychnine HCl | Sigma Aldrich | Cat# S-8753 |
| QX314 chloride | Abcam Cambridge, MA | Cat# ab120118 |
| CGP 55845 hydrochloride | Abcam Cambridge, MA | Cat# ab120337 |
| THIP hydrochloride (gaboxadol) | Abcam Cambridge, MA | Cat# ab120426 |
| RNAscope fluorescent kit | ACD, Cambridge, MA | Cat# 320850 |
| RNAscope Gabrd-Mm-C3 probe | ACD, Cambridge, MA | Cat# 459481-C3 |
| RNAscope tdTomato- probe | ACD, Cambridge, MA | Cat# 317041 |
| Anti-GABA<sub>A</sub> $\delta$ Antibody, rabbit | Alomone labs, Jerusalem, Israel | Cat# AGA-014 |
| Anti-rabbit Alexa 647 | Abcam, Cambridge, MA | Cat# 150083 |
| Anti-rabbit Alexa 488 | Abcam Cambridge, MA | Cat# 150081 |
| rabbit anti-GABA<sub>A</sub> receptor-α1 | Abcam | Cat# ab33299 RRID: AB_732498 |
| rabbit anti-GABA<sub>A</sub> receptor-α6 | GeneTex | Cat# GTX130947 |
| mouse anti-GABA<sub>A</sub> receptor-β3 | Neuromab | Cat# 75-149 RRID: AB_2109585 |
| rabbit anti-GABA receptor-γ2 | Synaptic Systems | Cat# 224 003, RRID: AB_2263066 |
| rabbit anti-GABA<sub>A</sub> receptor-δ 1:2000 | gift from Dr. W. Sieghart and Dr. Petra Scholze, Medical University of Vienna, Austria | |
| **Experimental models: Organisms/ Strains** |  |  |
| *flox*ed Gabrd mice (*Gabrd<sup>tm1.1Jmag/J</sup>*) | Dr. Jamie Maguire, Tufts University Jackson Labs | Stock# 023836 Lee et al., 2014 |
| Gabrd knock out mice (*Gabrd<sup>tm1Geh/J</sup>*) | Dr. Gregg Homanics, University of Pittsburgh Jackson labs | Stock# 003725 Mihalek et al., 1999 |
| Gabra6-cre mice B6.D2-Tg(Gabra6-cre)B1Lfr/Mmucd | MMRC | RRID: MMRRRC_015966-UCD Funschilling et al, 2002 |
| Floxed tdTomato reporter line (Ai14, Gt(ROSA)26Sor<sup>tm14(CAG-tdTomato)Hze/J</sup>) | Jackson Labs | Stock# 007908 Madisen et al., 2010 |
| **Software and Algorithms** |  |  |
| Igor Pro 6 | Wavemetrics | https://www.wavemetrics.com/ |
| MafPC | Courtesy of M.A. Xu-Friedman | https://www.xufriedman.org/mafpc |
| MATLAB (R2017a) | MathWorks | https://www.mathworks.com/ |
| Fiji (ImageJ) | NIH | https://fiji.sc/ |
| AxographX 1.7.0 | Axograph | https://axograph.com/ |
| Prism 6 | Graphpad | https://www.graphpad.com/ |
| Adobe Illustrator | Adobe | https://www.adobe.com/products/illustrator.html |
| Moseq2 | Sandeep Datta Laboratory | https://github.com/orgs/dattalab/teams/moseq2-users |
| Python | Python | https://www.python.org/ |

## CONTACT FOR REAGENT AND RESOURCE SHARING

Further information and requests for resources and reagents should be directed to and will be fulfilled by the Lead Contact, Wade Regehr.

## EXPERIMENTAL MODEL AND SUBJECT DETAILS

### Mice

Animal procedures have been carried out in accordance with the NIH and Animal Care and Use committee (IACUC) guidelines, and protocols approved by the Harvard Medical Area Standing Committee on Animals. *Flox*ed Gabrd mice (*Gabrd^tm1.1Jmag^*/J, Jackson Labs stock# 023836) and Gabrd knock-out mice (*Gabrd^tm1Geh^*/J Jackson labs stock# 003725) were obtained from Dr. Jamie Maguire (Tufts University). Gabra6-Cre (B6.D2-Tg(Gabra6-cre)B1Lfr/Mmucd) mice were obtained from the MMRC. A reporter line expressing tdTomato in cre-positive cells (*floxed* tdTomato line A14) was obtained from Jackson labs (stock# 007908). Animals were kept on a mixed background (129Sv/SvJ and B6/C57). Mice were housed under standard conditions in groups of 2-5 animals on a 12 h light-dark cycle with food and water available *ad libitum.* For parental behavior assays dams were singly housed. Adult animals of either sex 2-5 months of age were used for all experiments, including FISH, immunohistochemistry, electrophysiology and behavioral testing.

## METHOD DETAILS

### Fluorescence In situ hybridization (FISH)

Animals were anesthetized with isoflurane and the brain was rapidly removed and frozen on dry ice before embedding in optimal cutting temperature (OCT) compound (Tissue-Tek). Tissue was cut on a cryostat (Microm HM500-CM) at a thickness of 20 µm and mounted on glass slides (Superfrost Plus, VWR, 48311-703). Fluorescent in-situ hybridization was performed according to the ACD-Bio RNAscope Multiplex Assay manual, (document Number 320513) with minor modifications. The samples were then fixed in 4% paraformaldehyde in phosphate-buffered saline (PBS) for 15 min at 4 °C, then dehydrated with 50% (×1), 70% (×1), and 100% (×2) ethanol washes for 5 min each. The slides were then air-dried and a barrier was drawn around the tissue section with an Immedge hydrophobic barrier pen (Vector Laboratories). The tissue was incubated in RNAscope protease III reagent (ACD-Bio 322337) at room temperature for 30 min, then rinsed twice in PBS for 5 min.

Fluorophore-conjugated probes, Gabrd-Mm-C3 probe (Cat# 459481-C3), tdTomato-C2 probe (Cat# 317041) were incubated with the slide-mounted tissue sections at 40 °C in a HybEz II oven (ACD-Bio) for 2 hours and washed twice in RNAscope wash buffer reagent (ACD-Bio 310091). Fluorescence amplification steps were then applied as follows: incubate in AMP 1-FL for 30 minutes at 40 °C (HybEZ oven), followed with 2x wash with 1x wash buffer for 3 min at room temperature, the tissue was then incubated in AMP 2-FL for 15 minutes at 40 °C, followed by 2x wash, then incubated in AMP 3-FL for 30 min at 40 °C, followed with 2x wash, and lastly was incubated in AMP 4-FL-A for 15 minutes at 40 degrees, then 2x wash. Sections were then stained with DAPI and mounted with ProLong antifade reagent (Thermo Fisher Scientific P36930). In addition, control probes were used to ensure the quality of the *in situ* experiment. The positive control is a cocktail of housekeeping genes (C1-Mm-Polr2a, C2-Mm PPIB and C3-Mm-UBC). The negative control probe targets bacterial RNA (C1, C2, C3-dapB). The slides were imaged using whole slide scanning microscope (Olympus VS120) with a 20X air objective.

### Immunohistochemistry

Mice were anesthetized with isoflurane and perfused with ice cold phosphate buffered saline (PBS, pH=7.4, Sigma Cat# P-3813), followed by a solution containing 4% paraformaldehyde in PBS. The brain was removed and postfixed in the same solution at 4 °C overnight. For slicing, the brain was embedded in 4% agar (Sea Plaque, Lonza, Cat# 50101) and then sliced in PBS using a vibratome (VT1000S, Leica) at a thickness of 50 µm. Antigen retrieval was performed prior to immunostaining. Slices were permeabilized in 0.2% TritonX (Sigma Cat# T9284) in PBS for 30 min and then incubated in a solution containing 0.001 % trypsin (Sigma Cat# T5266) and 0.001 % Ca_2_Cl in PBS for 1 minute. Slices were then rinsed 3 times for 5 minutes in PBS and then blocked in a solution containing 4% normal goat serum (NGS), 0.1% TritonX in PBS for 1 hour. After blocking, slices were incubated in the same solution with the addition of primary antibody and 0.001 % trypsin inhibitor (Sigma Cat# T6522) overnight at 4°C. Slices were then washed 3 times for 10 minutes an incubated in 4% NGS, 0.1% TritonX in PBS with addition of secondary antibody for 2 hours at room temperature. Slices were then washed 3 times for 5 minutes in PBS, mounted on glass slides (Superfrost Plus, VWR, Cat# 48311-703) and covered with mounting medium (ProLongDiamond, Thermo Fisher Scientific, Cat# P36961) and a glass coverslip. The mounting medium was allowed to cure for at least 24 hours before imaging.

### Imaging and image analysis

Whole-brain images were taken on an Olympus VS120 slide scanner, and confocal stacks were acquired on an Olympus FV1000 confocal microscope. Images were processed using standard routines in Fiji (ImageJ).

### Quantitative western blotting

Western blotting was performed according to standard protocols. For quantitative assessment of protein levels in mice brain tissues, fluorescently tagged secondary antibodies were used. Prior to harvesting tissue mice where anesthetized with isoflurane and perfused transcardially with ice cold PBS. The brain was then rapidly removed from the skull and tissue was dissected in PBS and frozen at -80°C until further processing. Tissue was lysed in a buffer containing 150 mM NaCl, 25 mM HEPES, 4 mM EGTA and protease inhibitor (Sigma-Aldrich, Cat# P8340). After addition of SDS harvested tissue underwent ten freeze-thaw cycles (−80 to +55 °C). After SDS-PAGE, gels were transferred onto nitrocellulose membranes and blocked with filtered 5% nonfat milk/5% goat serum in Tris-buffered saline for one hour at room temperature and incubated with primary antibodies with 5% BSA in Tris-buffered saline containing 0.1% Tween-20 (TBST) overnight at 4°C. Each membrane was incubated with primary antibodies against GABA_A_ receptor subunits as follows: rabbit anti-α1GABA_A_R(1:10 000; Abcam, Cat# ab33299, RRID: AB_732498), rabbit anti-α6GABA_A_R (1:1000; GeneTex, Cat# GTX130947), mouse anti-β3GABA_A_R (Neuromab, Cat# 75-149, RRID: AB_2109585), rabbit anti-γ2 GABA_A_R (1:1000; Synaptic Systems, Cat# 224 003, RRID: AB_2263066), rabbit anti-δGABA_A_R (1:2000; custom made, gift from Dr. W. Sieghart and Dr. P. Scholze, Medical University of Vienna, Vienna, Austria). The following fluorescent secondary antibodies were used prior to washing with TBST: donkey anti-mouse IRDye 800CW IgG (1:10,000; LI-COR, Cat. No.: 926-32212, RRID: AB_621847), donkey anti-rabbit IRDye 800CW IgG (1:10,000; LI-COR, Cat. No.:926-32213, RRID: AB_621848), donkey anti-mouse IRDye 680RD IgG (1:10,000; LI-COR, Cat. No.:926-32222, RRID: AB_621844), and donkey anti-rabbit IRDye 680RD IgG (1:10,000; LI-COR, Cat. No.:926-32223, RRID: AB_621845). Visualization was carried out with the LI-COR Odyssey® fluorescent scanner and software (LI-COR Biosciences). Blots were imaged using an Odyssey Infrared Imaging System Scan, at a resolution of 42 µm. Images were analyzed in NIH ImageJ Software.

### Slice preparation for electrophysiology

Mice to be used for electrophysiological recordings were retrieved from the animal facility and allowed to acclimate for at least 8 hours before the experiment to reduce stress. Animals of either sex aged 2-4 months were anesthetized in their home cage by introduction of an isoflurane-soaked cloth and then perfused transcardially under continued isoflurane anesthesia with ice cold cutting solution containing (in mM) 110 CholineCl, 7 MgCl_2_, 2.5 KCl, 1.25 NaH_2_PO_4_, 0.5 CaCl_2_, 25 Glucose, 11.5 Na-ascorbate, 3 Na-pyruvate, 25 NaHCO_3_, 0.003 (R)-CPP, equilibrated with 95% O_2_ and 5% CO_2_. The brain was rapidly dissected and the cerebellum was cut into 250-270 µm thick parasagittal slices in the same solution on a vibratome (VT1200S, Leica). Slices were then transferred to 34°C warm artificial cerebrospinal fluid (ACSF) containing (in mM) 125 NaCl, 26 NaHCO_3_, 1.25 NaH_2_PO_4_, 2.5 KCl, 1 MgCl_2_, 1.5 CaCl_2_, and 25 glucose, equilibrated with 95% O_2_ and 5% CO_2_ and incubated for 30 min. Slices were then stored at room temperature until recording for up to 6 hours.

### Electrophysiology

Whole-cell recordings were obtained from visually identified granule cells using a 40x water-immersion objective on an upright microscope (Olympus BX51WI). Pipettes were pulled from BF150-86-10 borosilicate glass (Sutter Instrument Co., Novato, CA) at resistances of 4-5 MΩ on a Sutter P-97 horizontal puller. Electrophysiological recordings were performed at ∼32°C. For voltage clamp recordings, the internal solution contained (in mM): 30 K-gluconate, 110 KCl, 10 HEPES, 0.5 EGTA, 3MgATP, 0.5 Na_3_GTP, 5 phosphocreatine-tris_2_, 5 phosphocreatine-Na_2_, 5 QX314 chloride. The chloride reversal potential was ∼ 0 mV. For current clamp recordings, the internal solution contained (in mM) 130 K-gluconate, 10 KCl, 10 HEPES, 0.5 EGTA, 3 MgATP, 0.5 Na_3_GTP, 5 phosphocreatine-tris_2_, 5 phosphocreatine-Na_2_. The chloride reversal potential was ∼ -65 mV, similar to what was reported for GCs (Brickley et al., 1996). Seal resistance for all GC recordings was >5 GΩ. Electrophysiology data were acquired using a Multiclamp 700B amplifier (Axon Instruments), digitized at 50 kHz, filtered at 4 kHz, and controlled by software custom written in IGOR Pro (Lake Oswego, OR). The recording ACSF included (in µM): 2.5 (R)-CPP, 5 NBQX hydrochloride, 2 CGP, 1 strychnine to block glutamatergic receptors, GABA_B_ receptors and glycine receptors, respectively. SR 95531 (100 µM) was used to block GABA_A_ receptor-mediated currents. The δGABA_A_R preferring agonist 4,5,6,7-tetrahydroisoxazolo[5,4-c]pyridin-3-ol (THIP, 1 µM) was used to activate tonic GABAergic currents. All drugs were purchased from Abcam (Cambridge, MA) or Tocris (Bristol, UK). Analysis of electrophysiological data was performed with custom routines written in IgorPro (Wavementrics, Lake Oswego, OR) or in AxoGraphX.

### Behavioral testing

All behavior testing was performed in adult mice (>8 weeks old) of either sex with the experimenter blind to the genotype. Before behavioral testing, mice were transferred to the behavior room and allowed to acclimate for at least 30 min. Animals were housed on a 12 h light-dark cycle, and the experiments were carried out at the beginning of the dark cycle. All equipment used for behavioral testing was cleaned with 70% ethanol in between experiments. Most behavioral testing was videotaped at ∼30 fps using a 720p USB Camera with IR LEDs (ELP, Ailipu Technology Co.) and the iSpy open source video surveillance software suite (ispyconnect.com). Off-line analysis was automated using custom scripts written in Matlab (Mathworks). Select behaviors were scored live (pup retrieval, olfactory testing), or analyses was carried out manually (grooming, forced swim).

### Gait analysis

To assess baseline locomotion, gait analysis was run over the course of 8 consecutive days with 5 trials done for each animal per day. A custom video recording setup was used for evaluating gait patterns in mice. The setup consisted of an infrared illuminated transparent linear corridor (64.5(L) x 4 (W) x 6 (H) cm) atop a borosilicate glass floor. The corridor was flanked by two mirrors on the side angled at 48 degrees to project the images of the side views toward a video camera (Bonito CL-400B/C 2320 x 700 pixels at 200 frames per second, Allied Vision, Exton, PA.) situated beneath the glass floor. Three views were captured simultaneously at 200 fps as the animal walk down the linear corridor in a self-initiated manner. To annotate the locations of the body parts (nose, base of the tail, tip of the tail, left/right forepaws, and left/right hind paws), a convolutional neural network based on the stacked hourglass-network (Newell et. al., 2016) was trained using PyTorch on 500 manually annotated sample frames. We used the neural network to annotate frame-by-frame which results in a time series for the location of all aforementioned body parts. A Hidden Markov Model (HMM) was applied on the time series to segment each video into individual gait cycle. For a single gait cycle, we measured the following parameters: cadence, the number of cycles per second in Hz; stride length, the maximum distance traveled by the forepaw within a single cycle; paw width, the lateral distance between the two diagonal supports at any given time; stance duration, the average amount of time that the paws are on the ground during; velocity, the average speed of the center of mass; and tail elevation, the absolute height of the tip of the tail from the floor. We used a linear mixed effect model of the form Y ∼ Intercept + genotype + sex + genotype:sex + (1|name) and ANOVA for statistical test in the non-covariate case. For the covariate analysis involving the velocity we fitted a model of the form Y ∼ Intercept + Velocity + genotype + sex + genotype:sex + velocity:genotype + velocity:sex + (1|name) and ANOVA for testing statistical significance.

### Rotarod

To evaluate motor learning, we performed a rotarod assay over the course of three consecutive days, with 5 trials performed each day. Mice were placed on a rotating rod device (Rotamex-5 Rota-Rod, Columbus Instruments, controlled by Rotamex-5 software) running at 4 rpm baseline speed. After brief habituation, acceleration of the rod was initiated. The rod accelerated from an initial speed of 4 rpm to a maximum speed of 40 rpm in 10 second intervals. An animal fall was detected by infrared photo cells crossing the space above the rod and the time-to-fall was recorded. A fall was recorded either if the mouse fell off the rod, or loops around the rod without running. Animals were allowed to rest for 1 min between trials.

### Eyeblink conditioning

Adult mice of either sex performed motor behavior experiments during the dark cycle when they are most active and alert. The behavior setup was housed within a ventilated, anechoic and sound-insulated behavior chamber. Prior to the experiment, animals were implanted with a head bracket and allowed to recover for four days of post-surgery. To accustom the animals to head-fixation and the treadmill, five days of habituation were performed prior to the first day of eyeblink training. During habituation, the animal was head-fixed atop of a motorized treadmill six inches in diameter rotating at 20 mm/s. The treadmill was kept at this speed throughout the habituation period. During training, a white LED flash was used as the conditioned stimulus (CS) and a 50 ms, 15 psi air puff directed at the opposite eye was delivered as the unconditioned stimulus (US). The air puff was delivered via a 21 gauge blunt tip needle that is mounted on a manipulator to allow for individual adjustments. The pneumatic and electronics necessary for the control of the air puff was based on the design of Openspritzer (Forman et al., 2017). To record the movement of the eyelid, we illuminated the behavior chamber using an IR lamp and recorded the eye with a high-speed camera (Mako U-029B, Allied Vision, Exton, PA.) and a macro lens (1/2” 4-12mm F/1.2, Tamron, Commack, NY.) at 300 fps. The vertical span of the eyelid opening is measured as a function over time using a custom MATLAB script. An inter-stimulus-interval of 500 millisecond was used. 100 trials of CS-US pairing and 10 trials of only the CS were presented to the animal per day for 15 days. The inter-trial interval was randomized between 4 to12 seconds.

### Open field

Animals were placed in an uncovered rectangular behavior arena (30.3 cm x 45.7 cm, 30.5 cm high) containing fresh bedding and observed for 10 min without intervention. Analysis was performed offline using Matlab (postion, velocity, path traveled).

### 3-chamber assay

Sociability was assayed with the 3-chamber task. The behavioral arena consisted of a clear rectangular Plexiglas box (40.5 cm wide, 60 cm long, 22 cm high) without a top cover and divided into 3 equally sized compartments by clear walls. Each divider contained a 10.2 cm x 5.4 cm rectangular opening to allow navigation between the compartments. The left and the right chamber contained inverted wire cups (10 cm in diameter). Before behavioral testing, mice were allowed to navigate the middle chamber for 5 minutes with openings to the adjacent chambers closed. During a 10-minute habituation session, the doors were then opened and mice were allowed to freely navigate all 3 chambers for 10 minutes in the absence of any stimuli. After habituation, the doors were closed again while the animal remained in the central chamber. A social stimulus (juvenile mouse aged 15-30 days of the same sex and strain) and a novel object (mouse-sized plastic toy, Schleich GmbH, Germany) were placed in opposite wire cups. The sides of social and non-social stimuli were randomly selected to control for preference to either side of the arena. After stimulus placement, the doors were opened and the animal was observed for another 10 minutes. Automated analysis was performed offline using Matlab.

### Light/dark chamber

A light-dark chamber assay was used to measure anxiety-like in adult mice. The experiment consisted of a Plexiglas arena with the same outer dimensions as used for the 3-chamber assay, but with different interior dimensions. The light-dark chamber arena consisted of two chambers, with one removable divider of the same dimensions as used in the 3-chamber assay, including the door which remained open throughout. The light chamber was 40.5 cm x 40 cm chamber and brightly lit (>600 lux) and uncovered. The dark chamber was 40.5 cm x 20 cm and covered with darkly tinted Plexiglass that allowed videotaping of the mouse with an IR camera. Light intensity inside the dark chamber was <10 lux. At the beginning of the experiment, the animal was placed in the dark chamber and was allowed to freely navigate both chambers for 10 min.

### Motion Sequencing

Motion Sequencing (MoSeq)-based behavioral analysis was performed as in Wiltschko et al, 2015 and Markowitz et al 2018. In brief, MoSeq uses unsupervised machine learning techniques to identify the number and content of behavioral syllables out of which mice compose their behavior; identifying these syllables allows each video frame of a mouse behavioral experiment to be assigned a label identifying which syllable is being expressed at any moment in time. Behavioral phenotypes that distinguish wild-type and mutant mice can be identified by comparing differences in how often individual syllables are used in a given experiment. Here, individual mice (n = 12 per genotype) were imaged for 1-3 60-minute-long sessions using a Kinect2 depth sensor while behaving in a circular open field under red light illumination. These 3d imaging data were submitted to the MoSeq pipeline, which includes mouse extraction, denoising, and alignment steps before computational modeling. As has been done previously (Wiltschko et al, 2015), the kappa parameter (which sets the timescale at which syllables are identified) was specified by matching the distribution of syllable durations to a model-free changepoint distribution. All mice were submitted simultaneously for joint modeling with a shared transition matrix. MoSeq identified 64 syllables that make up 90 percent of the frames that comprise the dataset; for convenience, in Figure 4, we depict genotype-driven differences in the top 30 used syllables, which account for approximately 70 percent of the frames that comprise the dataset. Significant differences in syllable expression between genotypes were determined via permutation testing for each matched syllable pair (10000 rounds of usage value reassignment across cohorts). Significant syllables whose mean expression differed at an alpha level of 0.05 (with Benjamini and Hochberg FDR correction for multiple comparisons) are labelled in Figure 4.

### Pup interaction assay and spontaneous retrieval

Naïve virgin females were placed in a behavioral arena (30 cm x 46 cm, 31 cm high) without a top containing fresh standard bedding and were allowed to explore the arena freely during a 10-minute habituation period. After habituation, 3 pups of the same mouse strain aged P1-P3 were placed on one side, and 3 pup-sized novel plastic objects were placed on the opposite side of the arena. Interactions of the female with the pups and novel objects were recorded for 10 minutes (as defined as a 2.5 cm radius around the nest or the center of the toys). After 10 minutes one pup was removed from the nest and placed in the center of the behavioral arena. If retrieval of the pup had not occurred within the observation period the trial was counted as a failure to retrieve and was terminated. Pups were promptly returned to their mothers after the experiment. Analysis was performed offline using Matlab.

### Postpartum pup retrieval

The dam was allowed to remain in her home cage and 3 pups were removed from the nest and placed in the center and 2 corners of the cage opposing the nest (see Fig 5A for schematic). The time to retrieve each pup to the nest was scored once a day over a time period of 3 consecutive days, starting on P0. Care was taken not to disturb the dam and her pups unnecessarily during this period.

### Forced swim

Mice were placed in a glass cylinder (height, diameter) filled with water at room temperature and observed for a 6-min period. Animals were then removed from the beaker, towel dried and returned to their home cage. Videos were scored offline. Time spent mobile (swimming, actively struggling) and immobile (floating, with front paws and at least one hind paw immobile) were scored.

### Olfactory testing

To test the mice’s ability to detect and discriminate odors we performed a simple olfaction task. Animals were introduced to a clean, covered cage without bedding, food or water. Three neutral, non-social odors (coconut, raspberry, banana), 2 social scents (female and male urine), as well as a water sample were pipetted onto a clean cotton swab and introduced consecutively to the cage at random order. Mice were allowed to explore each scent for 5 min, and time spent investigating (sniffing, biting or chewing) was recorded live by the investigator.

## QUANTIFICATION AND STATISTICAL ANALYSIS

Electrophysiology data analysis was performed in Igor Pro (Wavemetrics), AxographX (Axograph) and Prism (Graphpad). The numbers of cells recorded are indicated in the figure legends, in the text and in a data table. To determine significance in a dataset the Whitney-Mann test, Wilcoxon signed rank test or Kruskal-Wallis test (with Dunn’s post-test) were used, as indicated. For select data sets, a one-sample t-test (Western blot analysis), or one-way ANOVA (gait analysis, with Dunnet’s multiple comparison post-test), were performed. Behavioral analysis was performed in Matlab (Mathworks) using custom written scripts, and Prism. Unpaired Student’s t-test was used to determine statistical significance. The number of animals used for each dataset is indicated in the text or figure legend.

## Supplementary Information

**Figure S1:**
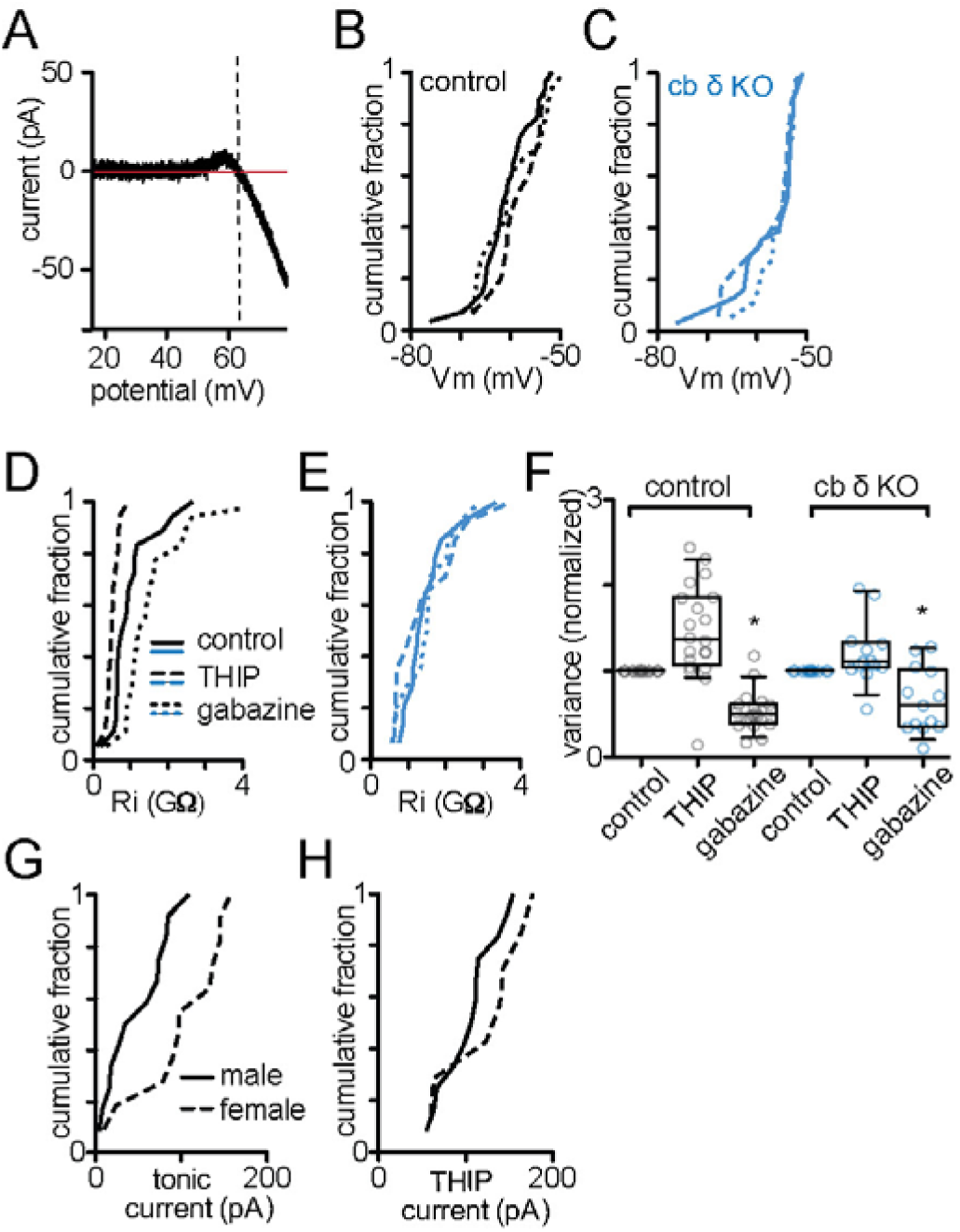
Electrical properties of control and cb δ KO granule cells (related to Figure 2) A) Example trace of cell-attached recording used to determine membrane potential (Vm), as described previously (Glickfeld et al., 2009). Graph shows current trace with the linear fit of the leak current (horizontal red line) subtracted. The vertical dashed line indicates the reversal potential of the current, corresponding to Vm. See methods for experimental details. B) Cumulative histograms of membrane potential (Vm) in control conditions (solid line), and in the presence of THIP (dashed line) or SR9931 (dotted line) for control animals. C) Same as (B) but for cb δ KO mice. THIP and SR95531 did not affect Vm in control or cb δ KO GCs (control: n=28 control solution, n=15 THIP, n=14 gabazine; cb δ KO: n=31 control solution, n=17 THIP, n=18 gabazine, (one-way ANOVA with Dunnett’s multiple comparison post-test). D) Cumulative histogram of input resistance (R_i_) in control conditions (solid line), or in the presence of THIP (dashed line) or SR995531 (dotted line). THIP decreased and SR95531 increased input resistance (R_i_) in control GCs (n=18 for all conditions, p<0.0001). E) In cb δ KO GCs THIP and SR95531 do not affect Ri (p>0.05). All data (D-E) describe matched observations, one-way ANOVA and Dunnett’s multiple comparison post-test. F) Box plot of normalized current variance in the presence of THIP or SR95531. THIP increased and SR95531 decreased current variance (measured as SD from mean current and normalized to control solution) in control granule cells (n=18 for all conditions, p<0.0001). In cb δ KO granule cells only SR95531 decreased current variance (p<0.05), while THIP has no significant effect (p>0.05; n=14 cells for all conditions). All data describes matched observations, one-way ANOVA and Dunnett’s multiple comparison post-test) G) Cumulative histograms of tonic current measured in males (solid line) and females (dashed line). Tonic current in females was higher than in males (males: n=12; females: n=11, p<0.01, KS test) H) Cumulative histogram of the THIP evoked current measured in males (solid line) and females (dashed line). The THIP evoked current is similar in males and females (males: n=12; females: n=11, p>0.1, KS test)

**Figure S2:**
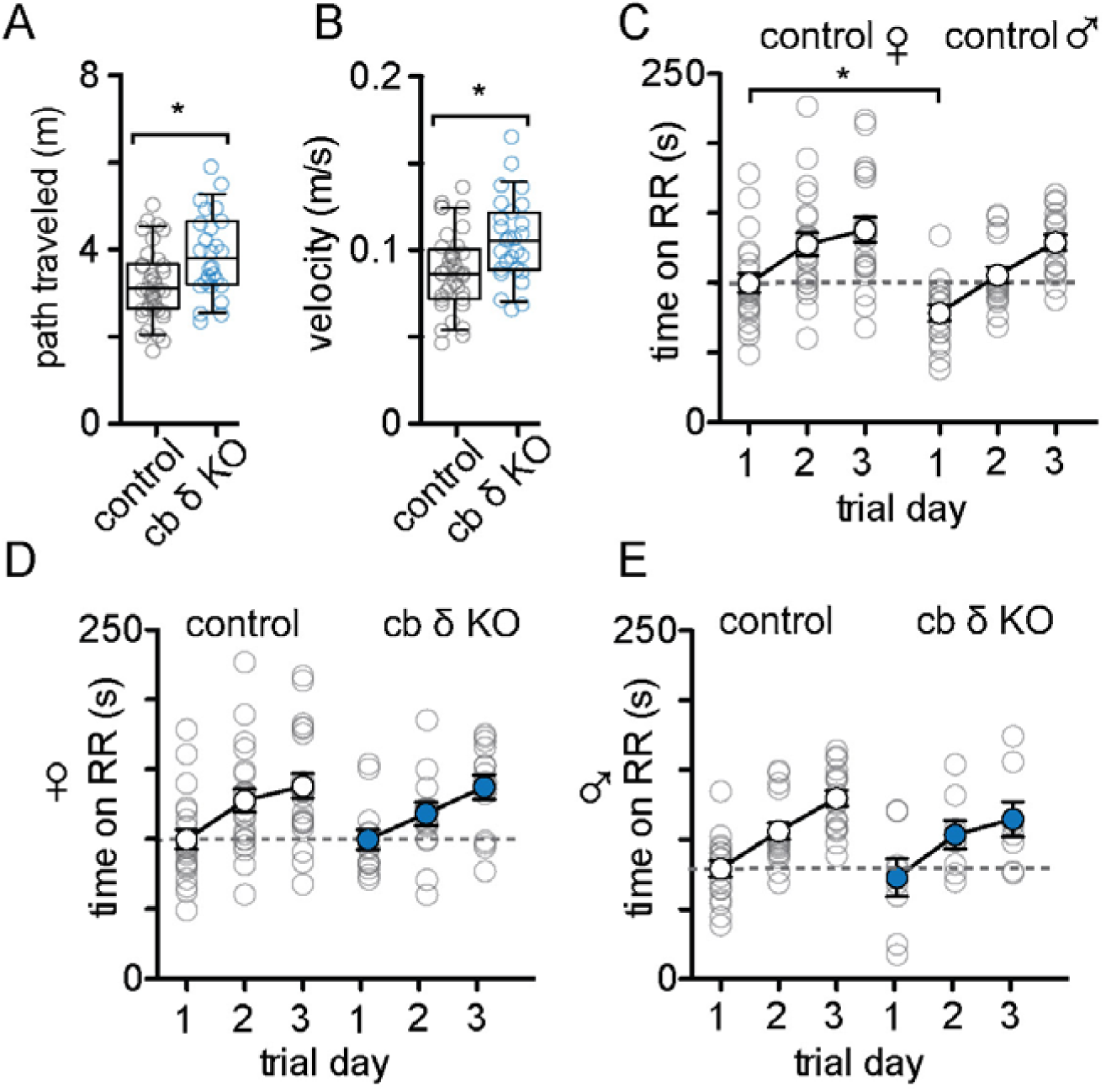
baseline locomotion and motor learning (related to Figure 3) A) The average path travelled during an open field behavioral task (control: n=37 animals, animals, cb δ KO: n=27 animals, p<0.007, Mann-Whitney test). Box represents interquartile range and median, whiskers are shown as 10-90 percentile. B) The average velocity during an open field behavioral task in control and cb δ KO animals (control: n=37 animals, cb δ KO: n=27 animals, p>0.005, Mann-Whitney test). Box represents interquartile range and median, whiskers are shown as 10-90 percentile. C) Rotarod performance in male and female control animals. Females perform slightly better on training day 1 (females n=21, males n=17, p<0.05, Mann-Whitney test) but not on consecutive training days. D) Rotarod performance of male control and cb δ KO animals (control n=17, cb δ KO n=8). E) Rotarod performance of female control and cb δ KO animals (control n=21, cb δ KO n=14).

**Figure S3:**
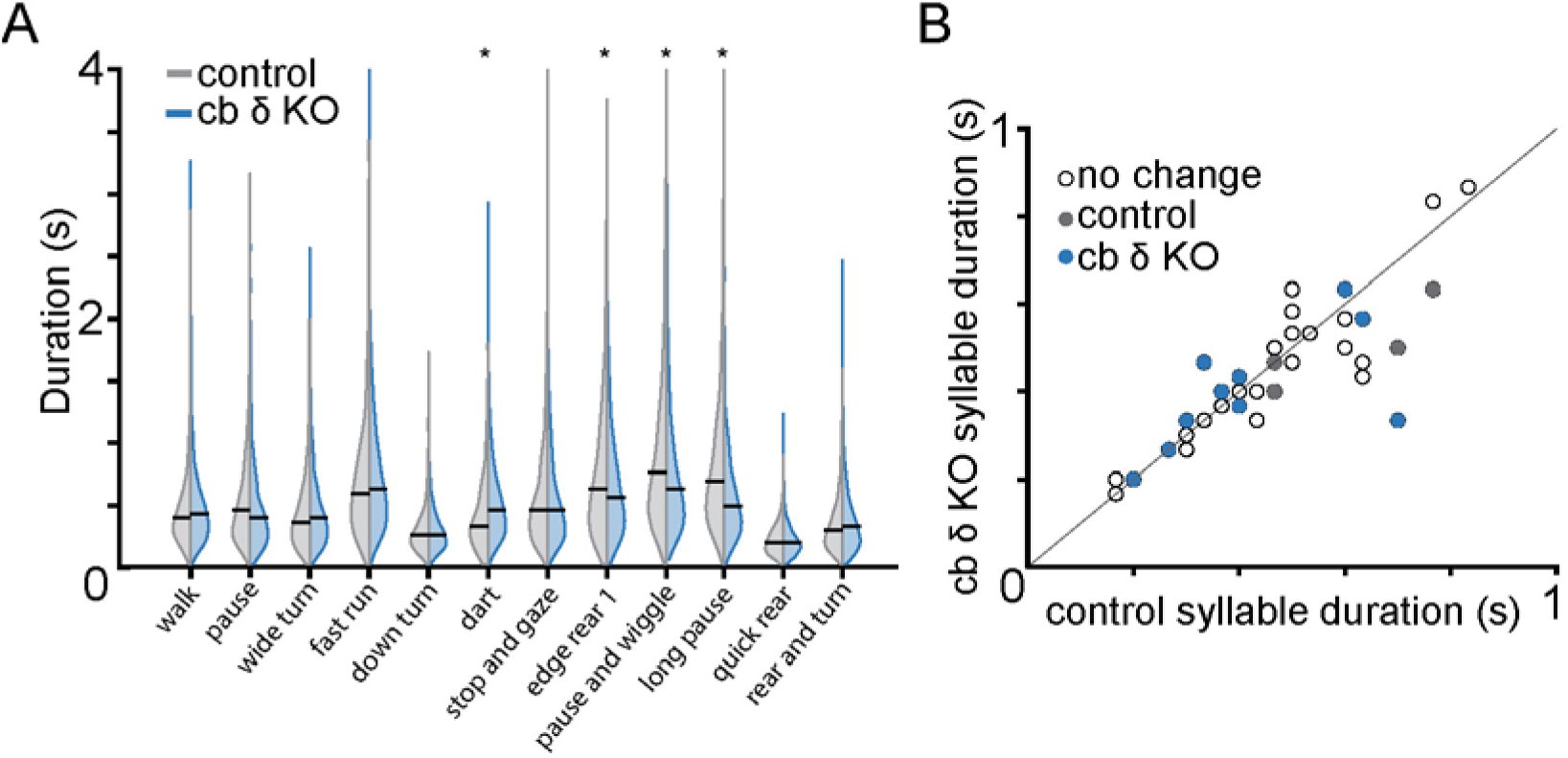
Syllable durations (related to Figure 4) A) Duration of select behavioral syllables for control (grey) and cb δ KO (blue) animals. B) Duration of select syllables performed by both control and cb δ KO animals plotted against each other. Syllables preferentially performed by control and cb δ KO animals are labeled in grey and blue, respectively. Clear circles denote non-regulated syllables. The duration of a subset of syllables differs between control and cb δ KO (*dart* in cb δ KO; *pause and wiggle* and *long pause* in control).

**Figure S4:**
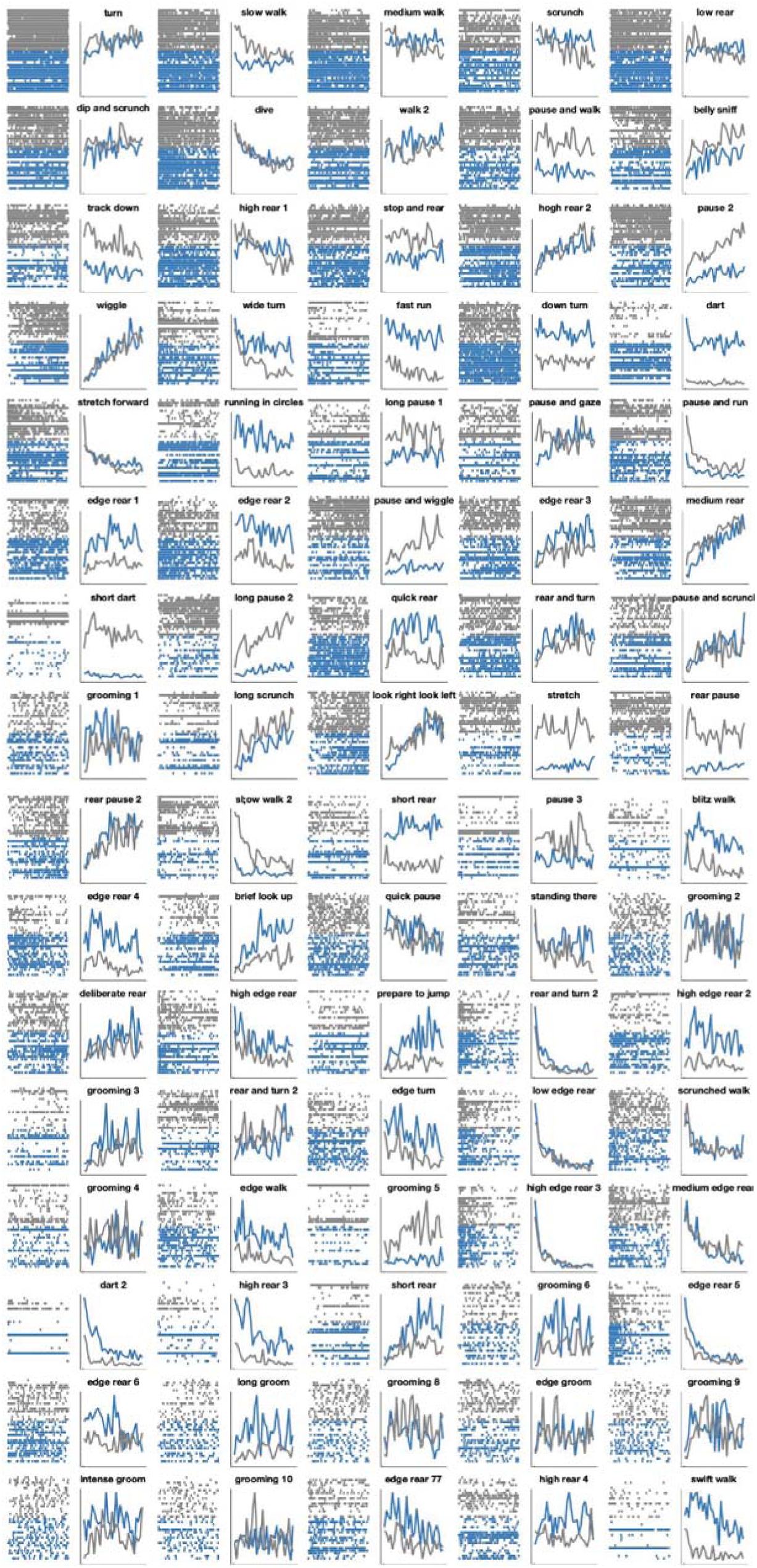
Behavioral syllables in the order of their frequency of occurrence (related to Figure 4) The most frequent syllables are shown with their respective IDs. Each panel shows syllable occurrence over the duration of the observation period in individual trials (left, tick plots) and median occurrence (right) in control (grey ticks and line) and cb δ KO (blue ticks and line).

**Figure S5:**
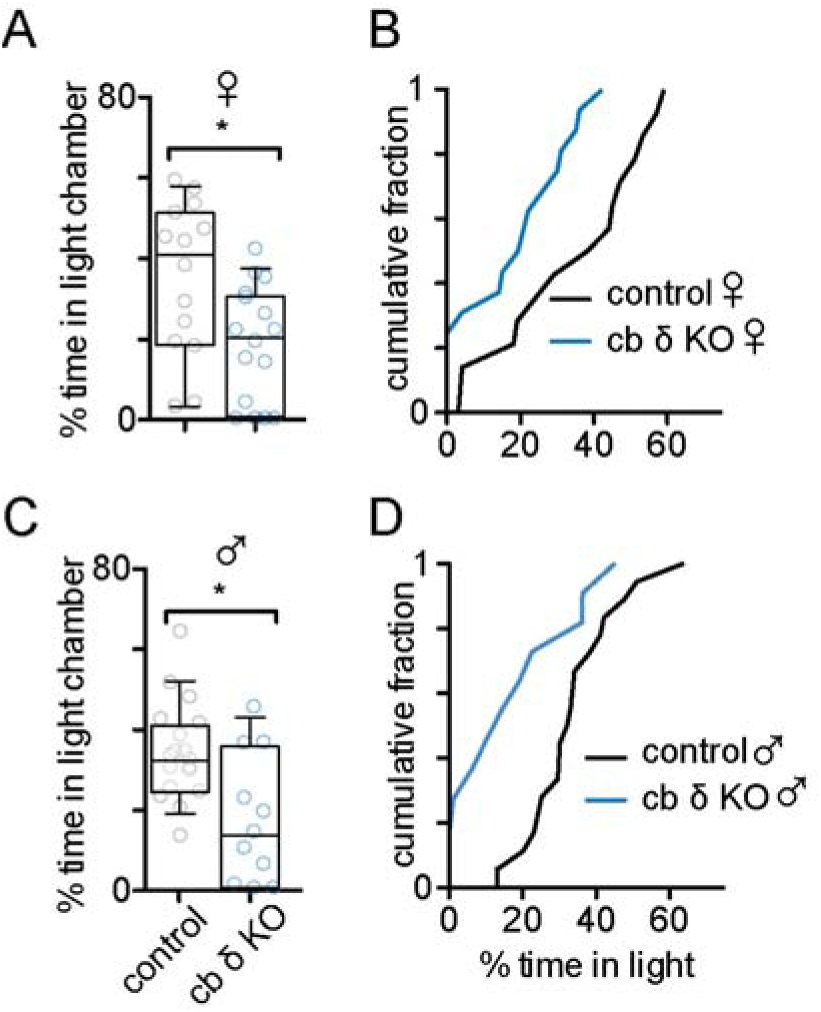
Both cb δ KO males and females display increased anxiety-like behavior (related to figure 5) A) Summary data of % time that females spent in the light compartment. Boxes denote interquartile range and median, whiskers represent 10-90 percentile. Circles show individual control (n=14, grey circles) and cb δ KO (n=16, blue circles) animals (p<0.03, Mann-Whitney test), B) Cumulative fraction of time spent in the light compartment (control: black line, cb δ KO: blue line) C) Summary data of % time that males spent in the light compartment. Circles show individual control (n=18, grey circles) and cb δ KO (n=11, blue circles) animals (p<0.05, Mann-Whitney test), D) Cumulative fraction of time spent in the light compartment (control: black line, cb δ KO: blue line)

**Figure S6:**
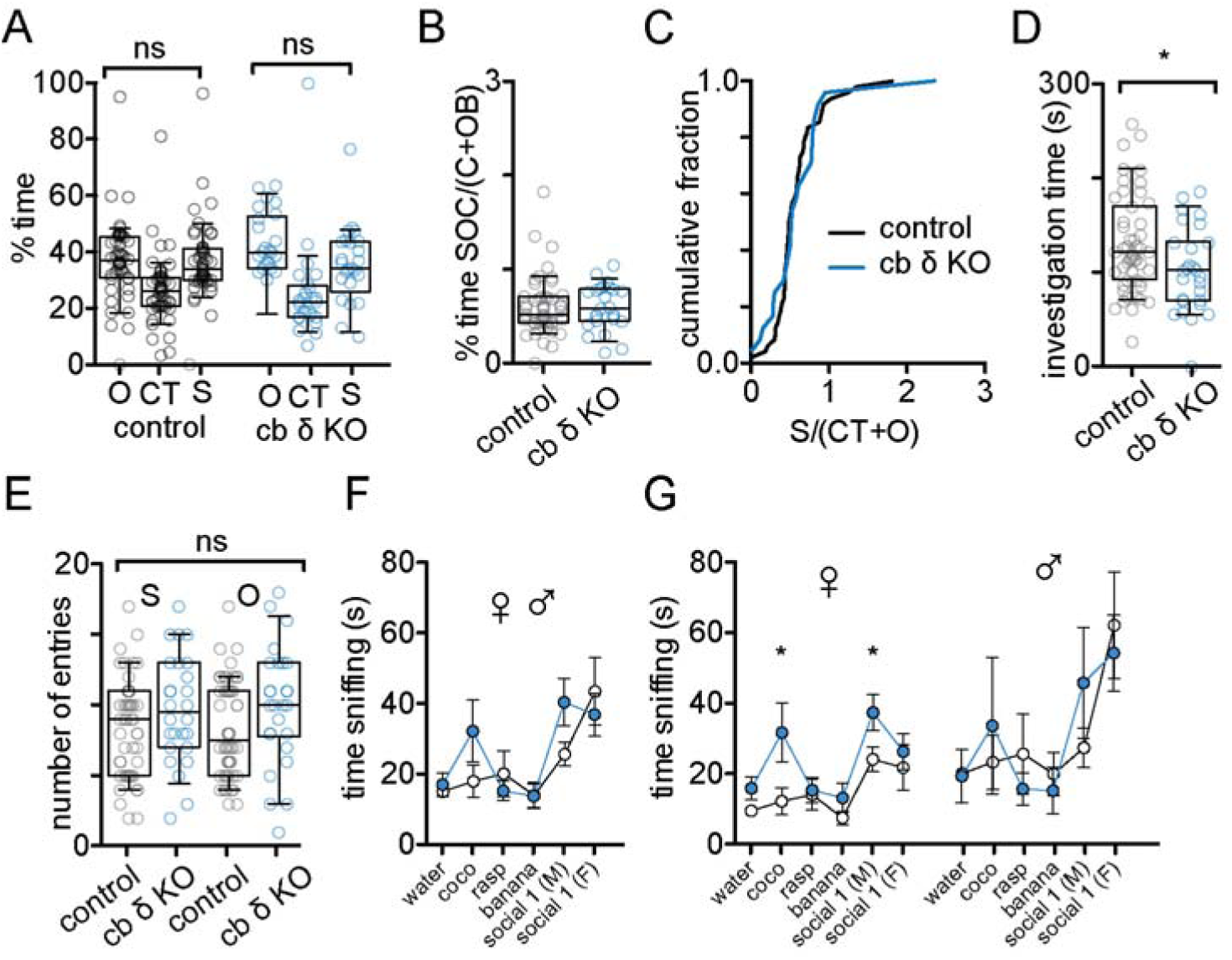
Baseline parameters for 3-chamber testing and olfaction (related to Figure 6) A) Summary data of 3-chamber assay under baseline conditions. In the absence of a social stimulus and an object stimulus neither control nor cb δ KO mice show a preference for either compartment of the 3-chamber arena. (control, n=55, p>0.2; cb δ KO, n=30, p>0.1, Wilcoxon signed rank test) B) In the absence of stimuli there is no difference in the S/(CTR+O) ratio between control and cb δ KO animals (control, n=55, cb δ KO, n=30, p>0.5, Mann-Whitney test) C) Cumulative probability of S/(CTR+O) ratios (p>0.2, KS test). D) Investigation time of social stimulus in control and cb δ KO animals (n=55, Mann-Whitney test, p<0.04) E) Control and cb δ KO animals enter the object and social compartments with similar frequency (control, n=50, cb δ KO, n=30, entries social chamber, p>0.2, entries object chamber, p>0.1, Mann-Whitney test) F) Average time spent sniffing four non-social (water, coconut, raspberry, banana) and two social cues (male and female urine) Control, n=24, cb δ KO, n=18, all odors, p>0.05, Mann-Whitney test. G) Increased time spent sniffing was noted in cb δ KO females for coconut and male odor (coconut, p<0.04; male urine, p<0.03, all other odors in females and all odors in males p>0.05, Mann-Whitney test)

**Figure S7:**
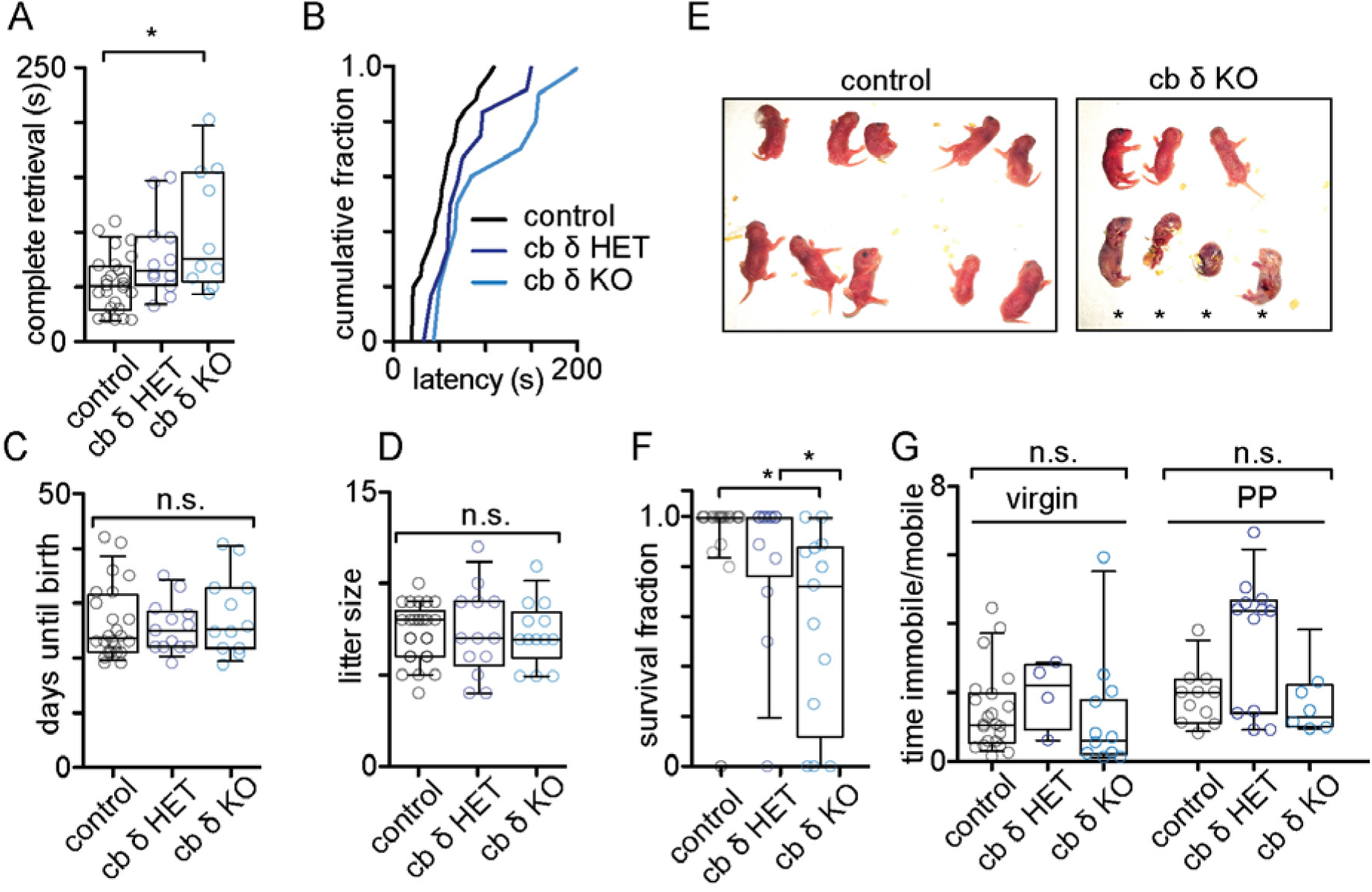
Retrieval, breeding postpartum and depression-like behavior (related to Figure 7) A) Time to complete retrieval of three pups to the nest for control, cb δ HET and cb δ KO dams. (p>0.02, Kruskal-Wallis test with Dunn’s post-test). B) Cumulative plot of complete retrieval time. C) The number of days from the first day of mating until birth are similar in control (grey circles, n=24), cb δ HET (dark blue circles, n=13) and cb δ KO (blue circles, n=11, p>0.6, Kruskal-Wallis test). D) Litter size at P0 is similar in control (grey circles, n=24), cb δ HET (dark blue circles, n=13) and cb δ KO (blue circles, n=11) females (Kruskal-Wallis test, p>0.9). E) Example litter of a control (left) and cb δ KO female (right). Approximately 12 h after birth, pups of a control dam are viable, cleaned and have nursed. Pups of cb δ KO females are often neglected (not cleaned, amniotic sac not removed) and/or cannibalized. Asterisks denote dead pups. F) Summary data of all control, cb δ HET and cb δ KO litter survival fractions. (p>0.0002; Kruskal-Wallis test with Dunn’s post-test). G) Ratio of time immobile and time mobile during the Porsolt forced swim test in virgin and postpartum (PP) females. (virgins: p<0.2; PP dams: p<0.07, Kruskal Wallis test with Dunn’s-post-test).

